# Diversifying stocking practices within rangelands supports diverse plant-pollinator communities and interactions

**DOI:** 10.64898/2026.05.22.727230

**Authors:** Avery E. Pearson, Laura Russo, John L. Neff, Braeden Duffee, Jada Martinez, Aidan McKinnis, Nick Medina, Julian Moore, Ethan W. Phillips, Elinor M. Lichtenberg

**Affiliations:** Department of Biological Sciences and Advanced Environmental Research Institute, University of North Texas, Denton, TX, USA; Department of Ecology and Evolutionary Biology, University of Tennessee, Knoxville, TN, USA; Central Texas Melittological Institute, Austin, TX, USA

**Keywords:** Coleoptera, Diptera, ecosystem services, Hymenoptera, Lepidoptera, rotational grazing, sustainability

## Abstract

Understanding how management alters species’ biodiversity and interactions is critical for sustainable agriculture. This can be especially important for rangelands, grasslands used for livestock grazing that cover 30% of land worldwide. In rangelands, ranchers need to maximize livestock production while supporting healthy ecosystems. Grazing management can alter the impact of livestock by varying grazing timing and intensity, but its ecological impacts remain poorly understood. One critical function that may be supported in these systems is pollination; how rangeland management may alter this ecosystem service is not systematically understood. We investigated how rotational stocking, where livestock are rotated among pastures, affected plant and pollinator communities and plant-pollinator interactions using working ranches in the Southern Great Plains. Sites were managed through continuous stocking or rotation with low or high frequency. We found that high-frequency livestock rotation supported greater flower abundance, and consequently greater pollinator abundance. Pollinator orders varied in their responses to management with Hymenoptera directly responding to management, and Coleoptera and Lepidoptera responding indirectly via changes in flower abundance. Community-level changes produced more specialized interaction networks that, along with species turnover, led to high interaction turnover across rangeland sites. This suggests that a mosaic of management strategies likely diversifies habitat and helps support plant and pollinator biodiversity at the landscape scale. This study demonstrates that rangelands can support plants, pollinators, and their interactions, particularly when managed with high-frequency rotational stocking.

## 1. Introduction

Rangelands – grasslands used for livestock grazing – exist at the intersection of agriculture and semi-natural habitat. They provide important habitat for both livestock and wildlife, covering 30% of land area globally and 35% within the U.S. (Bigelow and Borchers, 2017; Millennium Ecosystem Assessment, 2005). Rangelands provide key ecosystem services including livestock production, tourism, hunting, soil conservation, and nutrient cycling (Sala et al., 2017). These services hinge on ecosystem functions like pollination, biocontrol, and seed dispersal that result from ecological interactions among plants and animals. Continuation of these essential functions requires rangeland management that both promotes livestock production and supports organisms and their interactions. We focus on how management impacts insect pollinators and their plant hosts such as forbs (herbaceous flowering plants). Forbs enhance the nutritive value of livestock diets, counter soil compaction, slow water runoff, increase soil infiltration, fix soil nitrogen for improved grass growth, and provide vegetation structural heterogeneity that supports wildlife (DeBano et al., 2016; Muir et al., 2025). Despite pollinators’ critical role in maintaining healthy and resilient rangelands, how rangeland management regimes alter plant and pollinator communities and plant-pollinator interactions, and thus rangeland ecological functioning, remains a key knowledge gap.

Rotational stocking, where livestock are rotated among pastures, is increasingly being implemented. It aims to increase sustainability of livestock production while meeting profitability goals (Byrnes et al., 2018; Harmel et al., 2021; Teague et al., 2011) by enabling plant regeneration and distributing livestock impacts across the entire landscape rather than in a small number of preferred locations (Augustine et al., 2023). Rotational stocking practices vary in the size and number of pastures that livestock are rotated among, how frequently livestock are rotated, and the duration of each grazing event within a pasture (Allen et al., 2011). There is disagreement as to whether rotation benefits livestock production and graminoids (Augustine et al., 2020; Briske et al., 2008; Ge et al., 2025), but some evidence indicates that periods of rest from grazing do increase plant biomass (Ge et al., 2025) and flowering forbs (Busenitz et al., 2025; Ravetto Enri et al., 2017; Zhao and Iwaasa, 2022). This suggests that pollinators can benefit from rotational stocking. The few studies investigating rotational stocking impacts on pollinators find that pollinator diversity, abundance, and health increase under rotational stocking (Busenitz et al., 2025; Ravetto Enri et al., 2017). However, these impacts vary across insect orders (Ravetto Enri et al., 2017).

Because ecosystem functions like pollination are the result of species interactions, studying management impacts on plant and pollinator biodiversity cannot depict how rangelands respond to management. Understanding, and directly measuring, interactions is especially important given that plant and pollinator biodiversity can be poor predictors of plant-pollinator interactions (Olito and Fox, 2015; Poisot et al., 2012). This is partly due to the fact that plants and pollinators range from specialists, which interact with only one to a few other species, to generalists, which interact with a broad range of species (Ollerton, 2017). More specialized species are limited to sites (and thus management practices) that support their specific interaction partners. More generalized species can “rewire” their interactions by shifting to another species when partners become locally rare or extinct (Hagen et al., 2012; Lázaro and Gómez-Martínez, 2022). This helps confer resilience to plant-pollinator interaction networks with fewer secondary extinctions due to partner loss (Bascompte and Scheffer, 2023; Kaiser-Bunbury et al., 2010). These shifts alter ecosystem functioning (Kaiser-Bunbury and Blüthgen, 2015) by dictating which plants receive sufficient pollination (Ferreira et al., 2013). Rewiring often occurs before species become locally extinct, so we cannot capture these structural shifts without directly observing interactions (Valiente-Banuet et al., 2015).

Bipartite networks are a tool to quantify and visualize interactions (Bascompte and Jordano, 2014). Network structure is typically characterized through properties such as nestedness and degree of specialization. Variation in these metrics sheds light on ecosystem functioning (Kaiser-Bunbury and Blüthgen, 2015), including how plant-pollinator interactions respond to management. Nestedness is the tendency for specialists to interact with a subset of the generalists. Increased nestedness creates redundancy that enhances persistence of species and their interactions in the face of disturbance as long as species are able to rewire interactions (Bascompte and Scheffer, 2023; Coux et al., 2016; Zhang et al., 2011). High network-level specialization indicates that many species in a network are highly dependent on one or only a few other species. Since specialists and specialized interactions are the first to disappear, such networks are more sensitive to disturbance (Aguirre and Junker, 2024; Jacquemin et al., 2020; Kaiser-Bunbury and Blüthgen, 2015).

More recently, changes in network constituent species and structure, or interaction turnover, have been identified as important for understanding how management alters interactions. For example, turnover analysis showed that habitat loss increased rewiring to produce less nested and specialized networks (Lázaro and Gómez-Martínez, 2022). Interaction turnover is driven by species turnover and interaction rewiring (Fründ, 2021). Interaction turnover driven by species turnover can indicate more specialized networks. It can alternately indicate high resource availability and thus no need to switch partners (Simanonok and Burkle, 2014). Interaction turnover predominantly driven by rewiring can indicate a more resilient network structure (Valdovinos, 2019), more disturbed systems that contain mainly generalists that are capable of rewiring (Lázaro and Gómez-Martínez, 2022), or high competition environments that force some species to switch partners (Trøjelsgaard et al., 2015).

Though the research is nascent, there are indications that grazing alters plant-pollinator interactions. This has been studied only by comparing grazed to ungrazed land, grazing intensity gradients, or different grazer species. Networks from grazed pastures had greater species richness and structural complexity than ungrazed lands (Oleques et al., 2019; Vanbergen et al., 2014; Welti and Joern, 2018), showing that rangelands can serve as important pollinator habitat. Grazing intensity comparisons do not yet provide a clear pattern, with network nestedness not responding to intensity (Lázaro et al., 2016) and other indicators of network complexity being highest at intermediate (Lázaro et al., 2016) or high (Yoshihara et al., 2008) grazing intensity. Because rotation can increase plant and pollinator diversity (e.g., Döbert et al., 2025; Ravetto Enri et al., 2017), it will also likely alter interaction networks.

We investigated how rotational stocking alters plant-pollinator interactions and how responses differed among insect taxa. We focused our investigation on ranches that were managed with continuous stocking, low-frequency rotation, or high-frequency rotation. Continuously-stocked ranches provided livestock with unrestricted access to one larger pasture for the entire growing season. Ranches using rotational stocking moved livestock among smaller pastures every few weeks to months (low-frequency rotation) or every few days (high-frequency rotation). We asked two questions that focused on community and network-level attributes, as well as species’ roles (Table 1). (1) Because community structure impacts network structure (reviewed in Vázquez et al., 2009), we first asked how rotational stocking alters flower and pollinator communities, and which plant or pollinator taxa respond to management. We hypothesized an indirect impact where more frequent rotation gives the forb community more time to regrow and reproduce, leading to more flowers and flowering plant species (Busenitz et al., 2025; Steffens et al., 2013) and thus a more abundant and diverse pollinator community (Hyjazie and Sargent, 2022; Kennedy et al., 2013; Kral-O’Brien et al., 2021; Sritongchuay et al., 2026). We also hypothesized that pollinating insect orders would differ in their responses to grazing because of their diverse resource requirements and disturbance tolerances (Lichtenberg et al., 2025; Rader et al., 2014). While both food and shelter resources likely drive responses to grazing, we focused on floral food and insect-flower interactions. (2) How does rotational stocking alter plant-pollinator interactions? We hypothesized that stocking practices that supported greater plant and pollinator diversity, and thus more available interaction partners, would have more specialized interaction networks (Junker et al., 2015). Under our hypothesized indirect impacts via the plant community, we expected highly-connected species (hubs) to mainly be plants (as in Olesen et al., 2007). Additionally, we hypothesized that plant and pollinator compositional differences across stocking practices would lead to high interaction turnover that was driven by species turnover rather than interaction rewiring (Trøjelsgaard et al., 2015).

**Table 1:** Questions we addressed and analyses we used to address each one.

| Question | Analytical approach |
| --- | --- |
| Question 1: How does rotational stocking alter flower and pollinator communities? |  |
| How does management alter attributes of plant and pollinator communities? | Structural equation model (SEM) testing impacts on pollinator abundance, directly and indirectly via flower abundance |
|  | SEM testing impacts on pollinator richness directly and indirectly via flower abundance |
|  | Linear regression testing impacts on flower richness |
| Which taxa respond to management? | Plant community composition analysis via multivariate analysis of abundance |
|  | Pollinator community composition analysis via multivariate analysis of abundance |
|  | Four SEMs testing impacts on abundance of each pollinator order |
|  | Four SEMs testing impacts on richness of each pollinator order |
| Question 2: How does rotational stocking alter plant-pollinator interactions? |  |
| How does management alter network-level attributes? | Linear regression testing impacts on nestedness (wNODF) |
| | Linear regression testing impacts on network-level specialization ($H_2'$) |
| What roles do individual taxa serve in the networks? | Hub analysis of site-level networks |
|  | Hub analysis of networks pooled by management strategy |
| Are network changes driven by species' identities or their interactions? | Turnover analysis (WN, OS, ST) |

## 2. Methods

### 2.1 Study area

This study took place in the Cross Timbers ecoregion of North Texas, in the Southern Great Plains. This region is characterized by tallgrass prairies intermixed with woodland. Rainfall is highly variable, ranging from 686-965 mm/year. Soil texture ranges from clay to loam (Griffith et al., 2007). We worked with landowners to establish 19 sites on working ranches across Cooke, Denton, and Wise counties (Fig.1A). Each site consisted mainly of uncultivated grassland not treated with herbicides, contained few shrubs and trees, and was relatively flat. Pastures the sites were in ranged from 8 to 240 ha. The study region is a highly agricultural landscape. The 1 km surrounding almost all sites was at least 80% semi-natural habitat, mainly rangeland. Two sites were surrounded by a mix of rangeland and cultivated crops: wheat, corn, and sorghum (Boryan et al., 2011; USGS, 2024). Site centers ranged from 1.5 to 29.3 km apart (median 10.6 km). Sites were managed in one of three ways (Fig. 1B). Six sites were managed under continuous stocking, where livestock were not rotated. Five sites were managed under low-frequency rotational stocking, where livestock were rotated into new pastures every few weeks. Low-frequency rotation and continuously stocked sites were grazed by cattle. Eight sites were managed under high-frequency rotational stocking, where cattle and sheep were rotated into new pastures every few days. Field sampling was conducted in the spring (April – early May), summer (late May – June), and fall (late September – October) of 2023 to coincide with the three distinct floral blooming periods in this region.

**Figure 1:**
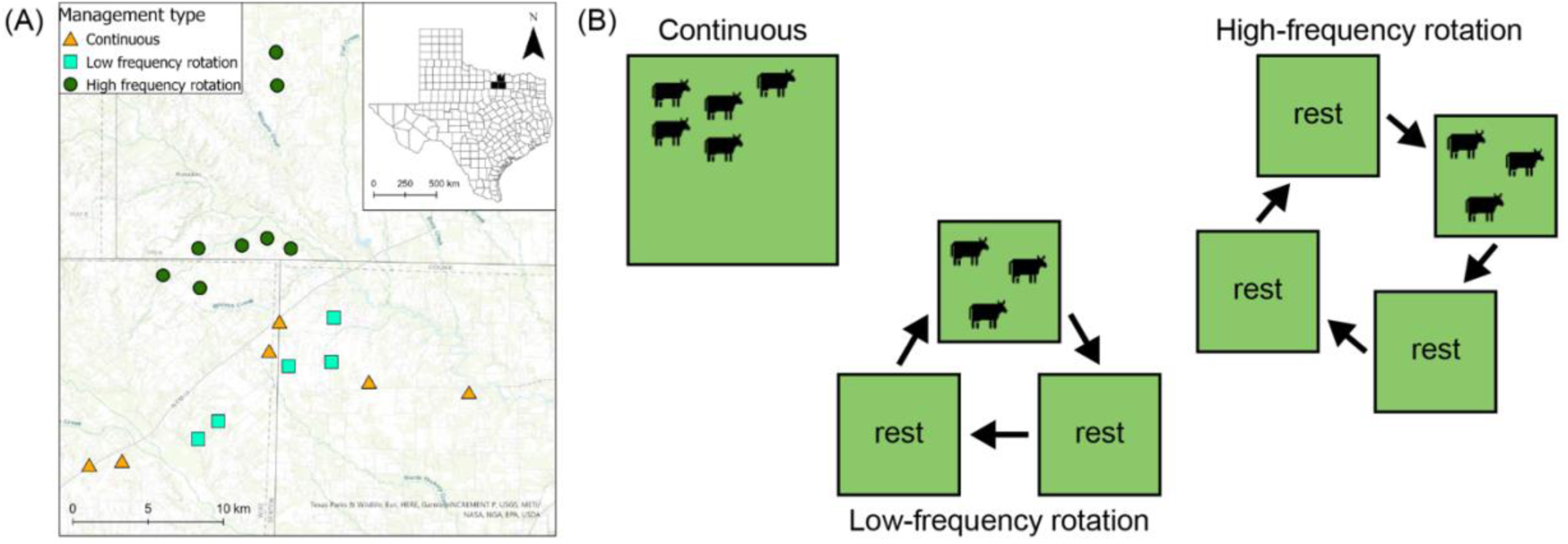
(A) Map of study site locations and (B) diagram of the grazing management strategies explored in this study. Sites under continuous stocking are represented by orange triangles, low-frequency rotation sites by teal squares, and high-frequency rotation sites by green circles.

### 2.2 Soil sampling

Since soil properties can influence plant and bee communities (Collins et al., 2025; Li et al., 2023) independent of management (Pennington et al., 2017; Renne et al., 2019), we measured soil texture as the percentage of sand, silt, and clay. We collected five soil samples per site: one in the center, and four 100 m from the center in each of the intercardinal directions (Fig. S1). We dug 15 cm deep holes and collected approximately 0.5 L of soil mixed from different depths. Soil texture was measured with a hydrometer using dry weights (Regen Ag Lab; Klute, 1986). We averaged across the five samples to get a sand, silt, and clay percentage for each site. Soil texture did not vary with management strategy (Kruskal-Wallis test: χ^2^_2_ = 0.98, *p* = 0.61; Fig. S2). Further analyses focused on soil sand content.

### 2.3 Flower sampling

To assess how grazing management strategies affected the flower community, we surveyed flowering plants along eight 90 m transects radiating from the center of each site in cardinal and intercardinal directions. This sampling area was 1% to 32% of the total pasture area. The proportion of the pasture sampled did not vary with management strategy (Kruskal-Wallis test: χ^2^_2_ = 2.60, *p* = 0.27). We placed 1×1 m quadrats every 15 m along each transect (48 quadrats per site; Fig. S1). We recorded the number and identity of any flowering angiosperms within each quadrat and counted the number of floral units (e.g., compound head, umbel, individual flower) to measure flower abundance (as in Lichtenberg et al., 2025). We also recorded the identity of all flowering angiosperm species encountered within the quadrats or within 1 m of each transect to more accurately measure flower richness. All flowering plant species were photo-vouchered and uploaded to iNaturalist for further identification (https://www.inaturalist.org/projects/lichtenberg-lab-rotational-grazing-study). 90% of plant taxa were identified to species, 2% to species group (closely related species that cannot reliably be distinguished), and 8% to genus (Table S1). We calculated each site’s total flowering forb abundance and richness across all quadrats and transects.

### 2.4 Pollinator sampling

To document the pollinator community and plant-pollinator interactions, we conducted aerial netting at each site. All netting sessions occurred between 09:00 and 16:00 in conditions that allowed normal pollinator activity: mostly to fully sunny weather, with temperatures above 18°C, and average wind speed under 6.7 m/s (Brittain et al., 2010; Buckles and Harmon-Threatt, 2019; Stein et al., 2020). We performed two 45-min netting sessions – one in the morning and one in the afternoon – at each site during each season. This totaled 114 aerial netting surveys, representing 85.5 hours of pollinator sampling. Sampling targeted flower visitors in the orders Coleoptera, Diptera, Hymenoptera (excluding ants), and Lepidoptera that were in contact with plant reproductive parts. We refer to these flower-visiting insects as “pollinators” even though flower visitation does not guarantee pollination. During netting sessions, one surveyor walked along and between the eight transects (Fig. S1) and captured all pollinators within 2 m of the transect. We recorded the flower species each captured insect was visiting. We recorded and released all captured monarch butterflies (*Danaus plexippus*) and bumble bee queens (genus *Bombus*) because these are vulnerable species (Davis et al., 2024; Hatfield et al., 2014). In the lab, we pinned and identified all pollinators to the lowest feasible taxonomic resolution and confirmed identifications with taxonomic experts (Table S2). 78% of taxa were identified to species or morphospecies, 2% to species group, 11% to genus, and 9% to family or subfamily (Table S3).

We used these data to calculate abundance and richness of all pollinators, and of each pollinator order, at each site, pooled across seasons in R (R Core Team, 2025). We estimated richness using iNEXT (Hsieh et al., 2016), which better accounts for taxa that potentially went undetected during sampling (Chao et al., 2005). We assumed that sites with no insects collected from a given order had a true richness of zero for that order; this assumption produced the same results as excluding those sites from relevant analyses. Richness calculations and network analyses excluded 22 damaged specimens (1% of the total) that could not be identified with sufficient taxonomic resolution to distinguish them as the same or different from other specimens that were identified to a finer resolution. Three of these specimens were Coleoptera, 13 Diptera, five Hymenoptera, and one Lepidoptera.

### 2.5 Plant-pollinator networks

We used the aerial netting data, which links pollinator species to the flowers they visit, to create quantitative bipartite plant-pollinator networks that documented the frequency with which each plant-pollinator taxon pair interacted (Memmott, 1999). Networks incorporated plants and insects at the same levels of taxonomic resolution as used to characterize each community. We pooled the morning and afternoon aerial netting data for each site across the spring, summer, and fall sampling events (Olsson et al., 2021). While there can be seasonal effects on plant and pollinator communities (Kimoto et al., 2012), pooled data produce larger networks that are better able to identify differences in interaction patterns across management strategies. This may have obscured seasonal responses to grazing management. Seasonal responses would best be identified by a multi-year study. Each network therefore represents six netting sessions, encompassing 270 minutes of surveying. We assembled networks using the bipartite package (Dormann et al., 2025).

For each network, we calculated nestedness (weighted NODF, or wNODF, which takes into account interaction frequencies; Almeida-Neto et al., 2008) and network specialization (*H_2_*’; Blüthgen et al., 2006). We chose these network-level attributes because they provide insight into network resilience, are common in the literature, and are robust to sampling effort (Rivera-Hutinel et al., 2012; Zoller et al., 2023). Moreover, wNODF incorporates interaction frequencies when evaluating nestedness, meaning it represents the importance of relative abundance, and *H′₂*is standardized and relatively robust to differences in network size.

We identified individual taxa that played important roles in networks via hub analysis. Hub analysis identifies species that are highly connected within and across a network (Dormann et al., 2025; Olesen et al., 2007). Such species have disproportionate impacts on the network’s structure (Burkle et al., 2021). This analysis determines each taxon’s within-module participation (*z* value) and among-module connectivity (*c* value). Plant and pollinator taxa with *z* > 2.5 and *c* > 0.62 are network hubs important for coherence of both a network and modules within the network. Taxa with high *z* but low *c* are module hubs that are important for coherence of the module they are in, while taxa with high *c* but low *z* are connectors that provide coherence across the network (Olesen et al., 2007). We ran this analysis separately for each site’s network, and for three networks that combined all sites and interactions within a management strategy.

### 2.6 Data analyses

To ensure that the spatial distribution of management strategies (Fig. 1A) did not strongly impact results, we tested for spatial autocorrelation. We assembled site by species abundance matrices for both plants and pollinators, excluding singletons to prepare data for the community composition analyses described below. We then calculated Bray-Curtis distances for the plant and pollinator communities (vegan package, Oksanen et al., 2025), and performed Mantel tests comparing community similarities (ade4 package, Dray and Dufour, 2007) to great circle distances among sites (sf package, Pebesma, 2018; Pebesma and Bivand, 2023) with 9,999 permutations. Neither plant (*r* = -0.09, *p* = 0.75) nor pollinator (*r* = -0.002, *p* = 0.49) communities showed spatial autocorrelation.

To determine how grazing management alters plant and pollinator communities (question 1), we used structural equation modeling (SEM). SEMs asked how management strategy (continuous stocking, low-frequency rotation, high-frequency rotation) and soil texture (as sand content) impacted pollinator abundance and richness both directly and indirectly via flower abundance (Fig. 2A). Preliminary analyses revealed that flower abundance and flower richness were highly correlated (Spearman’s rank correlation: S = 120.2, *r*=0.9, *p*<0.0001), and flower abundance better estimated resource availability for pollinators. We then used piecewise SEM (Lefcheck et al., 2025), which allows for non-linear relationships among variables (e.g., negative binomial for overdispersed abundances) and the sample sizes that are typical of many ecology field studies but insufficient for traditional SEM (Lefcheck, 2016). Our SEMs combined general(ized) linear regressions with flower abundance, pollinator abundance, or pollinator richness as the response variable. Management and soil texture were predictors in all models, and flower abundance was a predictor in the pollinator models. We used negative binomial regression for abundance models and the identity link for richness models. To identify differences in flower or pollinator abundance among management strategies, we assessed overlap of the confidence limits of each management strategy’s coefficients (J. Lefcheck, pers. comm.). To more fully understand plant community responses to management, we additionally regressed flower richness on management strategy and soil texture and assessed differences among management strategies using a post-hoc Tukey test (modelbased package; Makowski et al., 2025). One taxon, *Cryptorhopalum* dermestid beetles, was highly abundant across all sites. Because this dominance could dwarf other patterns, and to assess the degree to which this taxon drove observed patterns, we also ran insect abundance SEMs excluding *Cryptorhopalum*.

**Figure 2:**
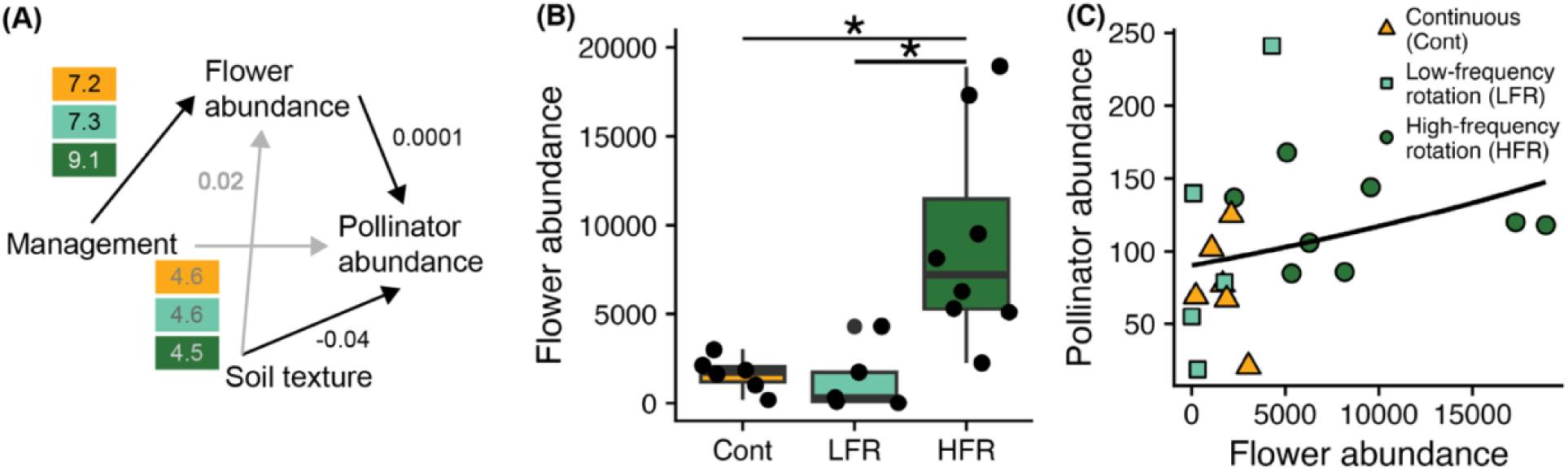
(A) Paths tested via structural equation modeling. Numbers are standardized path coefficients, with management coefficients listed top to bottom as continuous stocking, low-frequency rotation, and high-frequency rotation. Significant (p < 0.05) relationships are indicated by black arrows and black/white coefficients; gray arrows and text show relationships that were included but not significant. (B) High-frequency rotation sites had higher flower abundance. Stars (*) indicate significant differences between pairs. (C) Sites with higher flower abundance had higher pollinator abundance, irrespective of management practice. The curve is the best fit curve.

We investigated which taxa respond to management by (a) assessing community composition changes and (b) determining how each pollinator order responded to management. We assessed plant and pollinator community composition differences via multivariate analysis of abundance (mvabund package, Wang et al., 2022). Models included the site by species abundance matrices described above as responses; management strategy and soil sand content as predictors for the plant community composition analysis; and management strategy, soil sand content, and flower abundance as predictors for the pollinator community composition analysis. We tested significance of each term on the whole community using Monte Carlo resampling and a likelihood ratio test, followed by post-hoc tests comparing each pair of management strategies. Post-hoc analysis controlled for multiple comparisons using Holm free stepdown adjusted *p*-values. We additionally assessed the impacts of management and soil texture on each species, adjusting for multiple comparisons using a step-down resampling procedure. To determine how different pollinator orders responded to grazing management, we re-ran our SEMs using only specimens from each pollinator order (Coleoptera, Diptera, Hymenoptera, and Lepidoptera; total four SEMs). Since some of the sites did not have pollinators from all four orders, we did not analyze management impacts on community composition of each separate order.

To assess changes in plant-pollinator interactions (question 2), we used a regression approach to test if management strategy predicted network structure (wNODF and *H_2_*’). Because these metrics depend partially on network properties such as size and connectivity distribution, we calculated wNODF and *H_2_*’ z-scores that compared each site’s observed network to 500 null models generated using the Patefield algorithm (Patefield, 1981) that randomizes interactions while preserving species’ relative abundance (as in Russo and Stout, 2023). These z-scores were the response variables in linear regressions with management strategy and soil sand content as predictors. All regressions met model assumptions.

Finally, we assessed whether network changes were driven by species’ identities (species turnover) or their interactions (interaction rewiring) (Noreika et al., 2019; Poisot et al., 2012) using the betalinkr function in the bipartite package (Dormann et al., 2025). We calculated interaction dissimilarity (WN), interaction turnover (OS), and species turnover (ST) for each pair of sites (Fründ, 2021). We applied these calculations to both observed and null models, then calculated WN, OS, and ST z-scores for each network pair. These z-scores were the response variables in general linear mixed models with management comparison and distance between sites as fixed effects and the two sites being compared as random effects (glmmTMB package, Brooks et al., 2017). Management comparisons included: continuous stocking vs. continuous stocking, continuous stocking vs. low-frequency rotation, continuous stocking vs. high-frequency rotation, low-vs. low-frequency rotation, low-vs. high-frequency rotation, and high-vs. high-frequency rotation. We used post-hoc planned contrasts to ask whether dissimilarity (1) was greater among than within management strategies, and (2) differed among cross-strategy comparisons (emmeans package, Lenth et al., 2025).

## 3. Results

We observed 89,425 total flowers during the transect surveys, representing 166 unique taxa from 44 families. An additional 11 plant species were identified while netting insects (Table S1). We collected 1,959 pollinators composed of Coleoptera (36 taxa; 1,018 individuals; 52% of total captured pollinators), Diptera (45 taxa; 454 individuals; 23% of total captured pollinators), Hymenoptera (66 taxa; 307 individuals; 16% of total captured pollinators), and Lepidoptera (25 taxa; 180 individuals; 9% of total captured pollinators) (Table S3). These 172 taxa encompassed 56 families and 111 genera. A third of captured pollinators were *Cryptorhopalum* dermestid beetles (663 individuals: 224 at continuously-stocked sites, 262 at sites with low-frequency rotation, and 177 at sites with high-frequency rotation). We analyzed 1,936 flower-pollinator interactions representing 75 plant species and 487 unique plant-pollinator pairs. *Cryptorhopalum* visited 36 (48%) of the plant species. Site-level networks ranged in size from 12 to 58 unique interactions, averaging 36.7 interactions. Soils were almost all clay or clay loam, with soil sand content ranging from 10.6% to 64.7% and averaging 22.1% (Fig. S2).

### 3.1 Plant community – impacts on community attributes and the taxa that responded to management

SEM indicated that flower abundance increased at high-frequency rotation sites (β = 9.1; Fig. 2AB; Table S4). Linear regression found that flower richness was also higher at high-frequency rotation sites (Fig. S3A). Continuous (β_flower abundance_ = 7.2) and low-frequency rotation (β_flower abundance_ = 7.3) sites did not differ from one another in flower abundance or richness. Management strategy also significantly impacted plant community composition (Table S5), with all three management strategies differing from each other (continuous stocking vs. low-frequency rotation χ^2^ = 149.0, *p* = 0.002, continuous stocking vs. high-frequency rotation χ^2^ = 193.2, *p* = 0.001, low-vs. high-frequency rotation χ^2^ = 149.7, *p* = 0.002). No individual plant species responded to management (Table S6).

Soil texture, measured by soil sand content, was also an important predictor of the plant community. Sandier soils supported higher plant richness (linear regression: regression coefficient = 0.44; Fig. S3B), but had no impact on flower abundance (SEM β = 0.02; Fig. 3A; Table S4). Community composition analysis revealed that Virginia pepperweed (*Lepidium virginicum*), big-headed rabbit-tobacco (*Diaperia prolifera*), Texas toothleaf (*Stillingia texana*), dotted gayfeather (*Liatris punctata*), and hairy vetch (*Vicia hirsuta*) were more abundant at sites with higher soil sand content (10.8 ≤ χ^2^ ≤ 14.1; Table S6).

**Figure 3:**
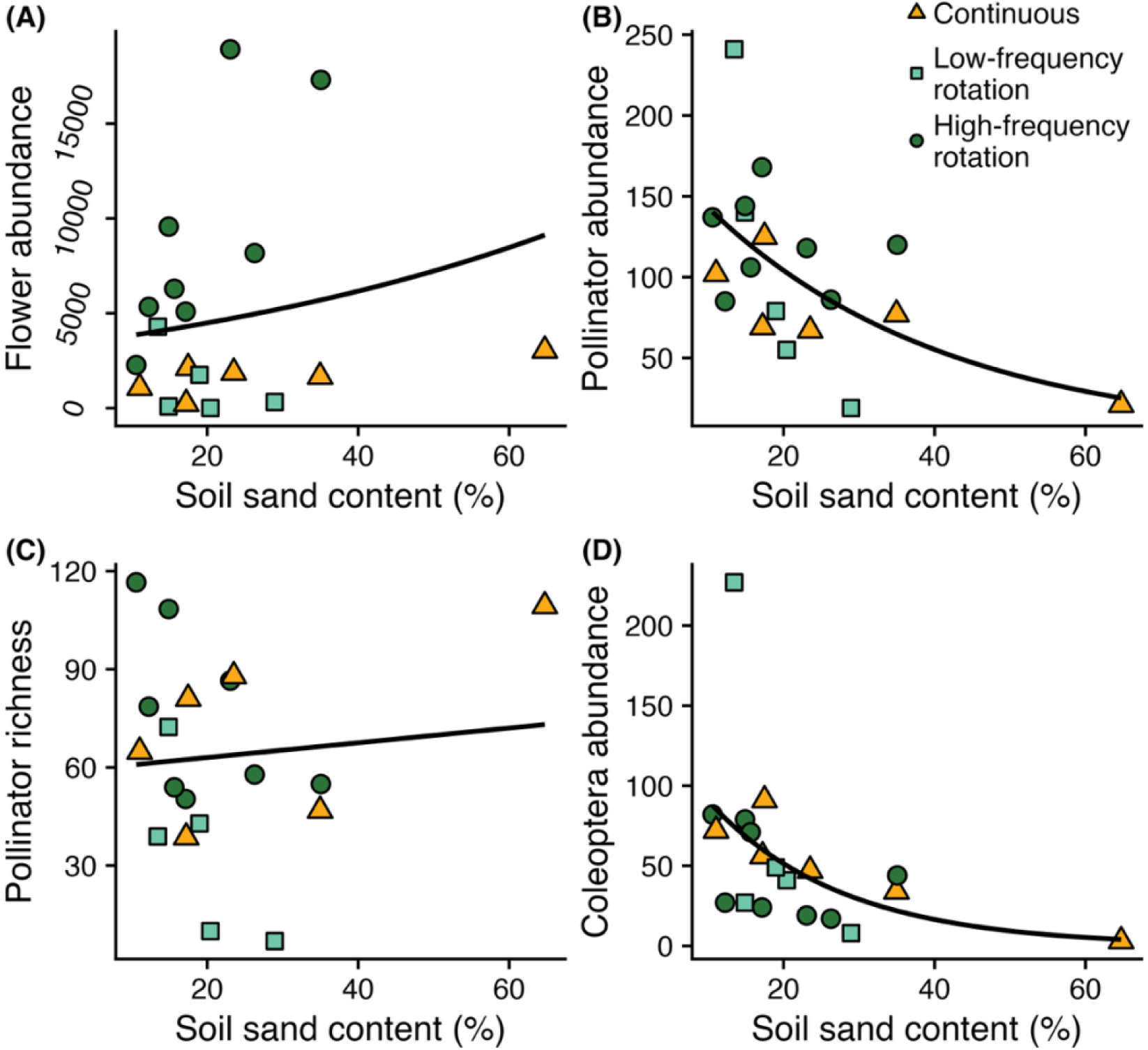
Impacts of soil texture on (A) flower abundance, (B) pollinator abundance, (C) pollinator richness, and (D) Coleoptera abundance. The curves and line indicate the best fit curves/line.

### 3.2 Pollinator community – impacts on community attributes and the taxa that responded to management

The pollinator community also responded to grazing management. SEM showed that management strategy indirectly, but not directly, drove pollinator abundance via its impact on flower abundance (Table S4). Specifically, higher flower abundance at high-frequency rotation sites supported increased pollinator abundance (β = 0.0001; Fig. 2A,C). However, impacts of flowers on pollinator abundance disappeared when the highly abundant *Cryptorhopalum* dermestids were excluded (β = 0; Table S4). Pollinator richness did not respond to management (SEM β_Cont_ = 70.1, β_LFR_ = 33.8, β_HFR_ = 77.1; Table S7). Pollinator community composition, on the other hand, did shift with management (Table S8) due to a difference between low- and high-frequency rotation sites (χ^2^ = 123.1, *p* = 0.006; continuous stocking vs. low- or high-frequency rotation χ^2^ = 103.4 and 100.0, respectively, *p* = 0.14 for both). As with the plants, no individual species responded to management (Table S9).

Soil texture also altered the pollinator community. Pollinator abundance increased in soils with less sand (SEM β = -0.04; Fig. 3B; Table S4), leading to shifts in the pollinator community composition (multivariate analysis of abundance; Table S5). However, this relationship disappeared when the *Cryptorhopalum* dermestids were excluded from the SEM (β = -0.02; Table S4), suggesting that their response to soil texture drove this relationship. The community composition analysis supported this, as *Cryptorhopalum* dermestids and another beetle – flower weevils (*Odontocorynus salebrosus*) – declined in abundance at sites with sandier soil (Table S6). Soil texture did not impact pollinator richness (SEM β = 0.1; Fig. 3C; Table S7).

SEMs that included just one pollinator order showed differences in whether insects responded to management, and whether they did so directly or indirectly via flower abundance (Figs. S4, S5; Tables S4, S7). An indirect impact, where sites with higher flower abundance supported more pollinators, was found for Coleoptera abundance (with and without *Cryptorhopalum*; β = 0.0001 for both), Lepidoptera abundance (β = 0.0001), and Lepidoptera richness (β = 0.0009). Grazing management directly increased Hymenoptera richness at sites experiencing high-compared to low-frequency rotation (β_HFR_ = 41.6, β_LFR_ = 3.6). Coleoptera abundance also responded to soil texture, decreasing at sites with less sand (with and without *Cryptorhopalum*; β = -0.08 and -0.05, respectively; Fig. 3D). Diptera and Hymenoptera abundance, and Coleoptera richness, did not respond to management strategy, soil texture, or flower abundance.

### 3.3 Plant-pollinator networks – network-level attributes

Interactions shifted across grazing management strategies (linear regressions; Fig. 4; Table S10). While nestedness z-scores did not differ (F_2,15_ = 2.6, *p* = 0.11), continuously-stocked sites were less specialized (i.e., lower *H_2_*’) than high-frequency rotation sites (F_2,15_ = 7.7, *p* = 0.005). Soil texture did not affect network structure (Table S10).

**Figure 4:**
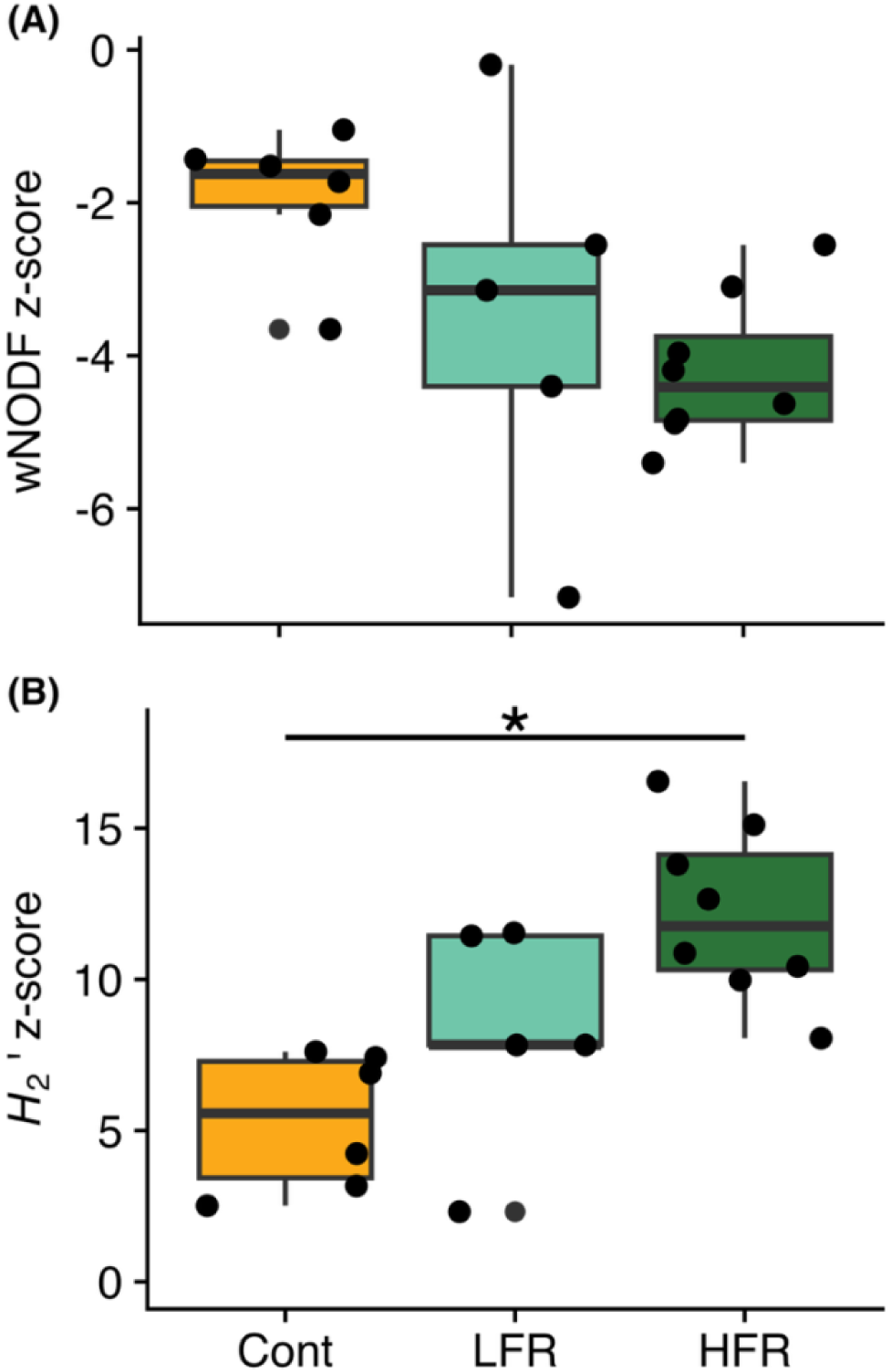
(A) Network nestedness (wNODF) did not change with management, but (B) network-level specialization (*H_2_*’) was lower at continuously-stocked sites compared to high-frequency rotation sites. “Cont” indicates continuously stocked, “LFR” indicates low-frequency rotation, and “HFR” indicates high-frequency rotation.

**Figure 5:**
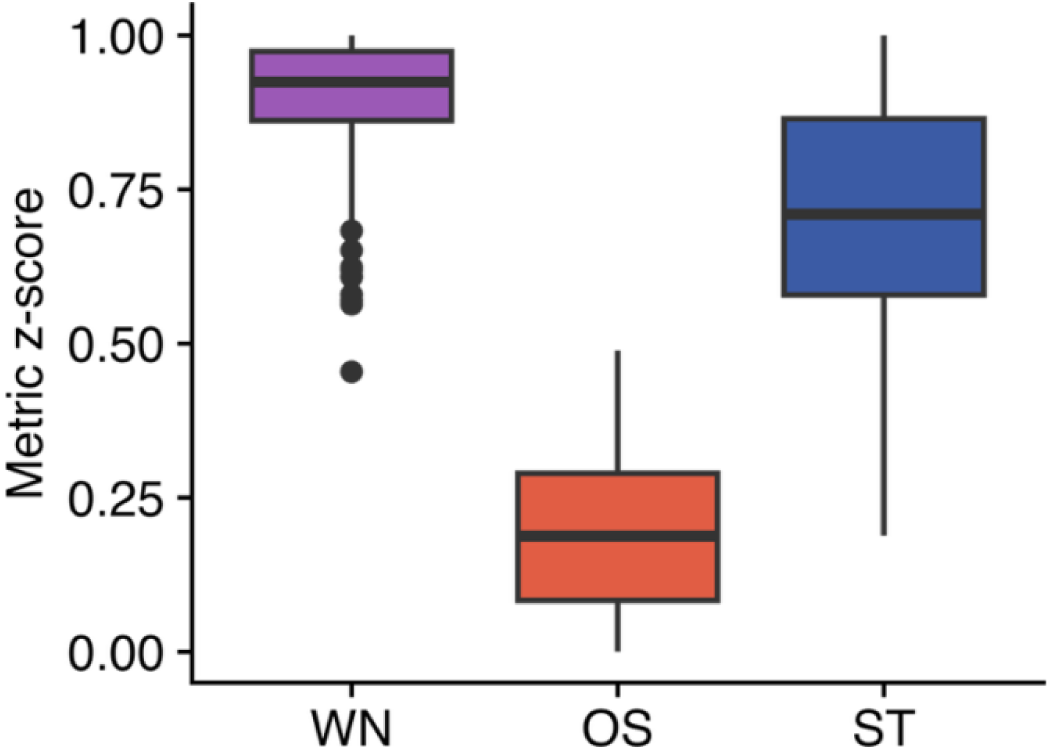
Interaction dissimilarity (WN), interaction turnover (OS), and species turnover (ST) between site pairs. Z-scores compared observed values to null models.

### 3.4 Network roles of plant and pollinator taxa

Hub analysis identifies taxa that are well-connected within or across network modules. Our analyses revealed that there were no plant or pollinator network hubs, that is, species that were both module hubs and connectors (Table S11). Plants served only as connectors, and not as module hubs. Insect taxa that served as hubs or connectors were evenly distributed across orders (Fisher’s exact test: *p* = 0.32). Diptera taxa were either only hubs or hubs and connectors, but never only connectors. Hymenoptera taxa were never both hub and connector, and Lepidoptera taxa were never only hubs. Insect (Fisher’s exact test: *p* < 0.0001) hub and connector taxa were more likely to be found in two or three seasons than expected by chance. At the order level, this pattern held for Diptera (*p* = 0.03), Hymenoptera (*p* = 0.001), and Lepidoptera (*p* = 0.03) but not Coleoptera (*p* = 0.08 with *Cryptorhopalum* and *p* = 0.13 without that genus). We did not see this pattern for plants (*p* = 0.19). Dark-winged fungus gnats (Coleoptera, Sciaridae; found in all three seasons), variegated fritillary butterflies (Lepidoptera, *Euptoieta claudia*; all three seasons), marginated calligrapher hover flies (Diptera, *Toxomerus marginatus*; all three seasons), honey bees (Hymenoptera, *Apis mellifera*; all three seasons), and flower weevils (Coleoptera, *Odontocorynus salebrosus*; spring and summer) emerged as module hubs at one site each and dermestids (Coleoptera, *Cryptorhopalum*; all three seasons) were module hubs at three sites. *Cryptorhopalum* were also connectors at three sites, while fleabane flowers (*Erigeron* sp.; spring and summer), marginated calligraphers, and blister beetles (Coleoptera, *Epicauta callosa*; summer and fall) were connectors at one site each. Table 2 shows hub and connector species from networks that aggregated sites to the management practice level. Overlap was low and occurred only between continuously stocked and high-frequency rotation sites.

**Table 2:** Hub and connector taxa in networks containing interactions pooled across all sites within a management practice. Text in parentheses provides ecological information. Plant taxa are identified with “P”, Coleoptera with “C”, Diptera with “D”, Hymenoptera with “H”, and Lepidoptera with “L”. We also indicate whether each taxon was found in the spring (“Sp”), summer (“Su”), or fall (“F”).

| Species role | Continuous grazing | Low-frequency rotation | High-frequency rotation |
| --- | --- | --- | --- |
| Module hub | Honey bees (H; Sp, Su, F)<br>Bee flies (*Geron*; D; Su, F)<br>Dark-winged fungus gnats (D; Sp, Su, F)<br>Flower longhorn beetles (*Typocerus octonotatus*; C; Su) | Flower weevils (C; Sp, Su) | Honey bees<br>Dark-winged fungus gnats<br>Flower longhorn beetles<br>Weevils (*Smicronyx*; C; F) |
| Connector | Desert chicory (*Pyrrhopappus pauciflorus*; P; Sp, Su)<br>Orange sulphur butterflies (*Colias eurytheme*; L; Sp, Su, F)<br>Spotted cucumber beetles (*Diabrotica undecimpunctata*; C; Sp, Su, F)<br>Sweat bees (*Lasioglossum disparile*; H; Sp, Su, F) | Gray hairstreak butterflies (*Strymon melinus*; L; Sp, Su, F) | American basketflower (*Plectocephalus americanus*; P; Su)<br>Lemon beebalm (*Monarda citriodora*; P; Su)<br>Orange sulfur butterflies<br>*Cryptorhopalum* (C; Sp, Su, F)<br>Spotted cucumber beetles<br>Feather-legged scoliid wasps (*Dielis plumipes*; H; Sp, Su)<br>Variegated fritillary butterflies (L; Sp, Su, F)<br>Bee flies<br>Convergent lady beetles (*Hippodamia* *convergens*; C; Sp,<br>Su)<br>Buckeye butterflies<br>(*Junonia coenia*; L;<br>Sp, Su, F)<br>Leafcutter bees<br>(*Megachile brevis*; H;<br>Su, F)<br>Northern paper wasps<br>(*Polistes fuscatus*; H;<br>Sp, Su, F) |

## 4. Discussion

Grazing management altered plant and pollinator communities, as well as plant-pollinator interactions. Specifically, plant abundance and richness increased at sites experiencing high-frequency livestock rotation. Pollinators showed mixed patterns, as pollinator richness did not respond to management while pollinator abundance showed an indirect response via a positive relationship with flower abundance. This largely reflected responses of Coleoptera and Lepidoptera. Community changes, in turn, altered the structure and identity of plant-pollinator interactions. While network nestedness did not differ across management strategies, high-frequency rotation sites showed higher specialization (*H_2_*’) than continuously-stocked sites. Taxa from all four pollinator orders served as module hubs or connectors, and plants served only as connectors. Insect hubs and connectors were more likely to be found in more than one season. There was high interaction dissimilarity across management strategies, due mainly to species turnover driven by distance among sites.

### 4.1. Rotational stocking impacts on flower and pollinator communities, and plant-pollinator interactions

The plant community response to management played a large role in altering pollinator communities and plant-pollinator interactions. Grazing management research mainly focuses on graminoids, but flowering plants provide vegetation with high nitrogen content that promotes livestock growth (Muir et al., 2025) and can comprise up to 20% of cattle diets (DeBano et al., 2016). As expected based on previous research (Briske, 2017), we found increased floral abundance and richness under high-frequency rotational stocking. This could be due to increased overall vegetation consumption that reduces grass-forb competition for sunlight (Koerner et al., 2018), reduced floral consumption as grasses are preferentially grazed (McNew et al., 2023) during a herd’s short time in a pasture, or sufficient rest for forbs to grow and reproduce (flower) under high-frequency rotation (Zhao and Iwaasa, 2022). Pollinators, on the other hand, responded to management mainly indirectly via changes to the flower community. This could have been driven by decreased competition with other pollinators (Doublet et al., 2022) or livestock (Black et al., 2011) as flowers became more abundant, or increased niche space as flower richness increased (Blüthgen and Klein, 2011). The low levels of nestedness (antinestedness) we observed suggests that pollinators did compete for flowers in our networks (Dormann et al., 2017) although antinestedness can also arise from sampling effects and other processes. Flower community differences also translated to more specialized networks at high-frequency rotation sites (as in Gómez-Martínez et al., 2022). This could be due to increased plant richness, or to changes in community composition such as retention of specialist or disturbance-sensitive species that were lost at continuously-stocked sites (reviewed in Soares et al., 2017). Flower richness and community composition, but not pollinator richness or community composition, differed between continuously-stocked and high-frequency rotation sites, further highlighting the important role that the flowering plant community played in structuring interactions. Despite this, plant species did not serve as module hubs (in contrast to, e.g., Dupont et al., 2014; Olesen et al., 2007). This indicates that pollinators, as well as plants, played important roles in structuring our networks.

### 4.2 Taxon-specific responses to management

Grazing management did not affect all orders of flower-visiting insects in the same way. This is consistent with a growing body of literature finding that pollinator responses to disturbance and changing resource availability vary widely (Aguirre-Gutiérrez et al., 2015; Bruninga-Socolar et al., 2022; Lichtenberg et al., 2025). Hymenoptera were the only group to respond directly to management. This may suggest that non-food resources such as shelter are important in structuring Hymenoptera communities (as in Collins et al., 2025). Coleoptera and Lepidoptera, on the other hand, only responded indirectly to management via their positive relationship with flowers. As Coleoptera were by far the most abundant pollinators in our study, it is unsurprising that they strongly influenced overall pollinator abundance responding indirectly to management and directly to soil texture (measured as relative sand content). Interestingly, Diptera were not sensitive to management or flower abundance. Diptera are commonly less sensitive than other pollinators to local habitat changes (Davis et al., 2023; Harrington et al., 2026), which could be partially because they are not central-place foragers and are thus less dependent on one site’s resources (Bergholz et al., 2022). A trait-based analysis that incorporates both floral food and shelter requirements may shed light on which attributes make pollinator taxa sensitive to grazing management. Potential management impacts on shelter for diverse pollinators include soil compaction, livestock waste that alters soil chemistry, and removal of herbaceous canopy or pithy stems.

*Cryptorhopalum* dermestids are abundant generalists that comprised nearly 1/3 of the total captured insects and visited almost half the plant species on which we documented pollinators. These beetles played important roles in the plant-pollinator interactions and community dynamics we observed. Removing this genus from the pollinator abundance SEM removed the negative impact of soil texture on pollinator abundance. This suggests that *Cryptorhopalum* are sensitive to soil texture, and potentially spend part of their life cycle under ground. Unfortunately, a thorough search including conversation with beetle experts, did not yield any *Cryptorhopalum* life history information. *Cryptorhopalum* also showed up as module hub and connector species, indicating they play important roles in structuring networks and connecting communities across landscapes (Olesen et al., 2007), thus promoting agroecosystem functioning (Allen et al., 2022). These roles are often seen for abundant generalist taxa (Aguirre and Junker, 2024; Olesen et al., 2007). While these beetles were highly abundant and structurally important, we commonly observed individuals sitting in flowers for extended time periods. Effective pollination requires pollen transfer and not just visitation (Pearson et al., 2023), and *Cryptorhopalum* may simply be consuming pollen (Evans, 2014). It is thus unclear whether these beetles can be functional substitutes for effective pollinators in disturbed landscapes.

### 4.3. Interaction turnover across sites

High interaction turnover across sites is common (Zhang et al., 2011), and in our system was driven by species turnover. High species turnover could indicate strong floral host preferences by pollinators (Simanonok and Burkle, 2014) or limited dispersal by our study organisms. Alternatively, it could represent environmental dissimilarity among sites, which altered local resource availability and thus the taxa a site could support (Maire et al., 2012; White et al., 2022). While not measured in this study, other work found that changes to local- and landscape-scale environmental variables altered the local pollinator community (e.g., Gámez-Virués et al., 2015; Sjödin et al., 2008). Our study region was fairly small, encompassed by a 16 km^2^ circle, but the impact of distance we found on interaction and species turnover would be consistent with an environmental gradient across the region. Further investigation into habitat impacts on plant and pollinator communities, from both taxonomic and traits-based perspectives, would provide insight into the ecological processes underlying our observed interaction turnover.

High species turnover in plant or pollinator communities may have led to the absence of a change to network nestedness that we observed. In other words, grazing management differences might have led to large compositional shifts rather than nonrandom losses of specific, specialized species. This conclusion is supported by the fact that community composition differed between low-and high-frequency rotation sites while network specialization did not. Other factors can also lead to the absence of a detectable change in nestedness. For example, if network hubs are lost, nestedness can remain the same while network resilience to perturbation decreases. Given the results of our other analyses, it seems more likely that different grazing management strategies lead to different network structures, rather than straightforward losses of specialized species.

### 4.4. Management and conservation implications

Our high-frequency rotation sites were grazed by a mix of cattle and sheep, while the other two strategies were implemented with just cattle. This reflects the fact that we worked on active ranches, a strength of our study that makes it more realistic than studies conducted solely on experimental ranches (Briske et al., 2011; Teague et al., 2011), and thus relied on the practices being employed by ranchers in our region. Sheep often preferentially graze forbs, and sheep grazing has been found to reduce forb abundance and diversity (Fornoff et al., 2025) and pollinators (Sharma Acharya et al., 2024), including when compared to cattle grazing (Cutter et al., 2022). Thus, the increased flower abundance and richness, and pollinator abundance, we found under high-frequency rotational stocking was likely due to rotation and not livestock species.

Working on active ranches and studying processes that happen at the landscape scale also necessitated that we conduct an observational study. Observational studies can include unrecognized influence by confounding variables and, as discussed above, greater variation than experimental studies that limit ability to make causal inferences (Dee et al., 2023). However, such studies also allow us to address questions that could otherwise not be studied (Siegel and Dee, 2025) and are integral to scaling from local ecological processes to landscape or ecosystem impacts (Gascuel-Odoux et al., 2022).

Our results indicate that rotation frequency is important. Our analysis of the taxa affected by management showed that plant and pollinator community composition differed between sites managed through low- and high-frequency rotation, and Hymenoptera richness was higher under high-than low-frequency rotation. Low-frequency rotation increases the likelihood that livestock graze a pasture for the entire bloom period of some plant species or that there is insufficient rest for plants to both regrow and reproduce (Steffens et al., 2013). Managing for pollinators thus should account for how rotation frequency and duration align with plant and pollinator phenologies (Black et al., 2011; Steffens et al., 2013). Interestingly, these community composition differences did not translate to differences in network attributes; network nestedness and specialization were the same at sites managed through low-and high-frequency rotation.

Because the ecosystem service of pollination relies on interactions among species, network level differences across our management strategies have implications for pollination and biodiversity conservation. The increased network specialization (*H_2_*’) at high-frequency rotation sites compared to continuous sites indicates that these networks contained plants visited by fewer pollinator taxa and pollinators that visited fewer plant species. Such networks often support higher functional diversity (Kühsel and Blüthgen, 2015). The larger number of connector taxa in the high-frequency rotation network may also help provide more consistent plant pollination services and pollinator floral resources (Burkle et al., 2021). Prolonged grazing pressure at continuously-stocked sites caused specialist species and interactions to be lost, and likely selected for disturbance-tolerant plants and pollinators, as seen in fragmented and simplified habitats (Gámez-Virués et al., 2015; Xiao et al., 2016). Plant species that were present at continuously-stocked but not high-frequency rotation sites tended to be introduced or noxious (*Buglossoides arvensis*, *Camelina microcarpa*, *Capsella bursa-pastoris*, *Carduus nutans*, *Cerastium* spp., *Erodium cicutarium*, *Hedypnois cretica*, *Helianthus annuus*, *Lactuca* spp.,

*Lamium amplexicaule*, *Melilotus officinalis*, *Rapistrum rugosum*, *Rumex pulcher*, *Stelaria media*, *Sisymbrium officinale*; Kartesz and BONAP, 2015) or common in disturbed habitats like roadsides (*Callirhoe involucrata*, *Camelina microcarpa*, *Capsella bursa-pastoris*, *Carduus nutans*, *Cnidoscolus texanus*, *Coreopsis lanceolata*, *Croton glandulosus*, *Erodium cicutarium*, *Hedypnois cretica*, *Helianthus annuus*, *Houstonia pusilla*, *Hybanthus verticillatus*, *Hypoxis hirsuta*, *Lamium amplexicaule*, *Lithospermum incisum*, *Melilotus officinalis*, *Nuttallanthus texanus*, *Oenothera laciniata*, *Physalis* spp., *Rapistrum rugosum*, *Rumex pulcher*, *Stellaria media*; Diggs et al., 1999). The very common *Cryptorhopalum* beetles were also more abundant at continuously-stocked sites. While continuously-stocked ranches may consist largely of disturbance-tolerant species (as happens with livestock forage under heavy grazing, Olff and Ritchie, 1998), they are likely still providing important pollinator habitat. A Species of Greatest Conservation Need (TPWD, 2025), *Megachile amica*, was found at one of our continuously-stocked sites, and connector and hub taxa overlapped between the continuous stocking and high-frequency rotation networks. Since community composition differed among grazing management strategies, a mosaic of management strategies likely diversifies habitat and helps support plant and pollinator biodiversity at the landscape scale.

Overall, our results demonstrate that rangelands can support speciose plant and pollinator communities with interaction networks that are less nested and more specialized than expected by chance. This was particularly true for sites managed with high-frequency rotation, which supported larger flowering plant and pollinator communities, and more specialized interactions. Plant and pollinator community composition varied among management strategies, and strategies had low overlap in hub and connector species. This suggests that each management strategy supports a distinct set of plant-pollinator interactions and there is no “one size fits all” best management approach. Rather, land managers need to consider their specific conservation goals and what is feasible for their operation. High-frequency rotation, for example, requires more fencing and labor to frequently move livestock to new pastures (Che et al., 2023). With such constraints, low-frequency rotation that is timed to permit flower growth in all pastures may achieve similar outcomes to high-frequency rotation. Interspersing management strategies across a landscape may also increase regional species pools and plant-pollinator interaction connectivity across pastures as long as more sensitive species are supported through management such as high-frequency rotation. However, such decisions need to consider whether the surrounding landscape contains more intensively-managed monoculture pastures, planted fields, or developed areas than our study region. Landscape composition differences may alter connectivity among pastures and thus differences among management strategies in plant-pollinator interactions. Seeding the plants we identified as connectors (desert chicory, American basketflower, lemon beebalm) may be another avenue for supporting plant-pollinator interactions (Monteiro et al., 2025). The low nestedness we found is similar to what is seen in newly-restored grasslands (Woodcock et al., 2026), indicating that there is potential to alter rangeland management in the Southern Great Plains to better support resilient interaction networks and thus ecological functioning. Improving understanding of how community composition and species interactions respond to management will help us better preserve biodiversity and ecosystem functioning in working landscapes (Glenny et al., 2022; Horak, 2014). Since rangelands cover ∼30% of land globally (Millennium Ecosystem Assessment, 2005), understanding how management can support plant-pollinator interactions offers huge potential to retain high quality habitat for pollinators and the ecosystem functions pollinators provide.

## Acknowledgements

We thank CP Cattle Company, the Dixon Water Foundation, Todd and Stephanie Underwood, and Wilson Land and Cattle for ranch access. Shannon Collins, Viktorya Dietrich, Marie Muñiz, Abbey Schedler, Laura Taylor, and Rob Whyle provided field and lab assistance. John Ascher, Val Bugh, Michael Caballero, Bill Dempwolf, Rick Fleischer, Jason Hansen, James McDermott, Hunter Messick, Sarah Paroski, Ed G. Riley, and Ashley J. Schmitz contributed to insect identification and verification. Two anonymous reviewers provided feedback. This material is based upon work that is supported by the National Institute of Food and Agriculture, U.S. Department of Agriculture, under award number 2022-38640-37488 through the Southern Sustainable Agriculture Research and Education program under subaward number OS23-162. USDA is an equal opportunity employer and service provider.

## Data availability

Data and code are available via Zenodo: https://doi.org/10.5281/zenodo.23192041.

