## Supplementary material for "Diversifying stocking practices within rangelands supports diverse plant-pollinator communities and interactions": Table S

**Table S1:** Abundance of each plant taxon we observed, and the management practice(s) under which it was found. Several species could not be reliably distinguished (e.g., doing so requires seeing fruit) and thus were identified to species group. We show both how many flowers we counted in transects and how many times we netted an insect visiting the plant taxon.

| Family | Taxon | Abundance in quadrats | # observed insect visits | Present at continuous sites? | Present at low frequency rotation sites? | Present at high frequency rotation sites? |
|---|---|---|---|---|---|---|
| Acanthaceae | *Dyschoriste linearis* | 20 | 0 | | | Y |
| Acanthaceae | *Ruellia* spp. | 0 | 0 | | | Y |
| Amaryllidaceae | *Allium* spp. | 1687 | 23 | Y | Y | Y |
| Amaryllidaceae | *Nothoscordum bivalve* | 162 | 27 | Y | Y | Y |
| Apiaceae | *Ammoselinum butleri* | 86 | 0 | Y | Y | |
| Apiaceae | *Ammoselinum popei* | 466 | 0 | | Y | |
| Apiaceae | *Bifora americana* | 14380 | 46 | Y | Y | Y |
| Apiaceae | *Chaerophyllum tainturieri* | 10657 | 1 | Y | Y | Y |
| Apiaceae | *Eryngium leavenworthii* | 91 | 43 | Y | | Y |
| Apiaceae | *Polytaenia texana* | 9 | 12 | | Y | Y |
| Apiaceae | *Torilis arvensis* | 2 | 1 | Y | Y | Y |
| Apiaceae | *Torilis nodosa* | 84 | 0 | Y | Y | Y |
| Apocynaceae | *Amsonia ciliata* | 301 | 15 | Y | | Y |
| Apocynaceae | *Apocynum cannabinum* | 227 | 3 | | | Y |
| Apocynaceae | *Asclepias asperula* | 7 | 54 | Y | Y | |
| Apocynaceae | *Asclepias viridis* | 15 | 427 | Y | Y | Y |
| Apocynaceae | *Matelea biflora* | 30 | 0 | Y | Y | Y |
| Asparagaceae | *Camassia scilloides* | 0 | 0 | | | Y |

| Family | Taxon | Abundance in quadrats | # observed insect visits | Present at continuous sites? | Present at low frequency rotation sites? | Present at high frequency rotation sites? |
|---|---|---|---|---|---|---|
| Asteraceae | *Achillea millefolium* | 33 | 1 | | | Y |
| Asteraceae | *Amphiachyris/ Gutierrezia* spp. | 8067 | 252 | Y | Y | Y |
| Asteraceae | *Arnoglossum plantagineum* | 0 | 0 | | | Y |
| Asteraceae | *Brickellia eupatorioides* | 31 | 0 | | | Y |
| Asteraceae | *Carduus nutans* | 0 | 0 | Y | | |
| Asteraceae | *Cirsium* spp. | 0 | 89 | Y | | Y |
| Asteraceae | *Coreopsis lanceolata* | 57 | 27 | Y | | |
| Asteraceae | *Diaperia prolifera* | 282 | 0 | | | Y |
| Asteraceae | *Diaperia verna* | 291 | 0 | | Y | Y |
| Asteraceae | *Engelmannia peristenia* | 74 | 11 | Y | | Y |
| Asteraceae | *Erigeron* spp. | 121 | 61 | Y | Y | Y |
| Asteraceae | *Gaillardia pulchella* | 257 | 30 | Y | Y | Y |
| Asteraceae | *Hedypnois cretica* | 2 | 0 | Y | | |
| Asteraceae | *Helianthus annuus* | 0 | 1 | Y | Y | |
| Asteraceae | *Helianthus maximiliani* | 0 | 7 | | | Y |
| Asteraceae | *Hymenopappus scabiosaeus* | 0 | 0 | | Y | Y |
| Asteraceae | *Lactuca* spp. | 4 | 0 | Y | Y | |
| Asteraceae | *Liatris punctata* | 27 | 29 | | Y | Y |
| Asteraceae | *Lindheimera texana* | 220 | 27 | Y | Y | Y |
| Asteraceae | *Marshallia caespitosa* | 0 | 0 | | | Y |
| Asteraceae | *Packera plattensis* | 13 | 8 | | Y | Y |
| Asteraceae | *Packera tampicana* | 122 | 45 | Y | Y | Y |
| Asteraceae | *Plectocephalus americanus* | 21 | 37 | Y | | Y |
| Asteraceae | *Pyrrhopappus pauciflorus* | 15 | 45 | Y | Y | Y |
| Asteraceae | *Ratibida columnifera* | 1 | 2 | | | Y |

| Family | Taxon | Abundance in quadrats | # observed insect visits | Present at continuous sites? | Present at low frequency rotation sites? | Present at high frequency rotation sites? |
|---|---|---|---|---|---|---|
| Asteraceae | *Rudbeckia amplexicaulis* | 221 | 1 | | | Y |
| Asteraceae | *Silphium albiflorum* | 0 | 0 | | | Y |
| Asteraceae | *Symphyotrichum ericoides* | 278 | 33 | | | Y |
| Asteraceae | *Symphyotrichum subulatum* var. *ligulatum* | 41 | 4 | | Y | Y |
| Asteraceae | *Tetraneuris linearifolia* | 3511 | 45 | Y | Y | Y |
| Asteraceae | *Thelesperma filifolium* | 228 | 23 | Y | Y | Y |
| Boraginaceae | *Buglossoides arvensis* | 0 | 1 | Y | | |
| Boraginaceae | *Lithospermum incisum* | 0 | 0 | Y | | |
| Brassicaceae | *Camelina microcarpa* | 0 | 0 | Y | | |
| Brassicaceae | *Capsella bursa-pastoris* | 2 | 0 | Y | | |
| Brassicaceae | *Lepidium virginicum* | 327 | 1 | Y | Y | Y |
| Brassicaceae | *Myagrum perfoliatum* | 0 | 0 | | Y | |
| Brassicaceae | *Physaria engelmannii* | 0 | 3 | Y | | |
| Brassicaceae | *Physaria gracilis* | 795 | 188 | Y | Y | Y |
| Brassicaceae | *Rapistrum rugosum* | 0 | 0 | Y | Y | |
| Brassicaceae | *Sisymbrium officinale* | 4 | 0 | Y | | |
| Campanulaceae | *Triodanis perfoliata* ssp. *biflora* | 0 | 0 | | Y | |
| Caprifoliaceae | *Symphoricarpos orbiculatus* | 0 | 0 | | | Y |
| Caryophyllaceae | *Arenaria serpyllifolia* | 225 | 0 | Y | Y | Y |
| Caryophyllaceae | *Cerastium* spp. | 0 | 0 | Y | | |
| Caryophyllaceae | *Paronychia virginica* | 7 | 0 | | Y | Y |
| Caryophyllaceae | *Stellaria media* | 3 | 0 | Y | | |
| Commelinaceae | *Commelina erecta* | 0 | 0 | Y | | |
| Commelinaceae | *Tradescantia ohiensis* | 0 | 4 | | | Y |

| **Family** | **Taxon** | **Abundance in quadrats** | **# observed insect visits** | **Present at continuous sites?** | **Present at low frequency rotation sites?** | **Present at high frequency rotation sites?** |
|---|---|---|---|---|---|---|
| Convolvulaceae | *Convolvulus equitans* | 1 | 0 | Y | | Y |
| Convolvulaceae | *Cuscuta* spp. | 617 | 0 | | | Y |
| Convolvulaceae | *Evolvulus nuttallianus* | 2 | 0 | | Y | Y |
| Euphorbiaceae | *Cnidoscolus texanus* | 0 | 0 | Y | | |
| Euphorbiaceae | *Croton capitatus* | 61 | 0 | Y | | Y |
| Euphorbiaceae | *Croton glandulosus* | 7 | 0 | Y | | |
| Euphorbiaceae | *Croton lindheimeri* | 5 | 0 | | | Y |
| Euphorbiaceae | *Croton monanthogynus* | 2885 | 3 | Y | Y | Y |
| Euphorbiaceae | *Ditaxis mercurialina* | 0 | 0 | Y | Y | Y |
| Euphorbiaceae | *Euphorbia bicolor* | 46 | 0 | Y | Y | Y |
| Euphorbiaceae | *Euphorbia davidii* | 1 | 0 | | Y | |
| Euphorbiaceae | *Euphorbia longicruris* | 1092 | 0 | Y | Y | Y |
| Euphorbiaceae | *Euphorbia missurica* | 162 | 1 | Y | Y | Y |
| Euphorbiaceae | *Euphorbia serpens* | 0 | 0 | | Y | |
| Euphorbiaceae | *Euphorbia spathulata* | 1158 | 6 | Y | Y | Y |
| Euphorbiaceae | *Stillingia texana* | 53 | 8 | Y | Y | Y |
| Euphorbiaceae | *Tragia* spp. | 681 | 0 | Y | Y | Y |
| Fabaceae | *Acacia angustissima* | 18 | 1 | | | Y |
| Fabaceae | *Astragalus crassicarpus/plattensis* | 0 | 0 | Y | | Y |
| Fabaceae | *Astragalus nuttallianus* | 314 | 0 | Y | Y | Y |
| Fabaceae | *Dalea aurea* | 2 | 0 | | | Y |
| Fabaceae | *Dalea compacta* | 6 | 1 | | | Y |
| Fabaceae | *Dalea purpurea* | 0 | 2 | | | Y |
| Fabaceae | *Desmanthus leptolobus* | 1 | 0 | | | Y |
| Fabaceae | *Lathyrus hirsutus* | 45 | 0 | | | Y |

| Family | Taxon | Abundance in quadrats | # observed insect visits | Present at continuous sites? | Present at low frequency rotation sites? | Present at high frequency rotation sites? |
|---|---|---|---|---|---|---|
| Fabaceae | *Medicago* spp. | 5778 | 7 | Y | Y | Y |
| Fabaceae | *Melilotus officinalis* | 0 | 0 | Y | | |
| Fabaceae | *Mimosa* spp. | 190 | 15 | Y | Y | Y |
| Fabaceae | *Neptunia lutea* | 0 | 0 | | | Y |
| Fabaceae | *Pediomelum cuspidatum* | 0 | 0 | | | Y |
| Fabaceae | *Pediomelum linearifolium* | 1 | 0 | | | Y |
| Fabaceae | *Senna roemeriana* | 0 | 0 | Y | Y | |
| Fabaceae | *Strophostyles leiosperma* | 0 | 0 | | | Y |
| Fabaceae | *Vicia hirsuta* | 277 | 0 | Y | | |
| Fabaceae | *Vicia ludoviciana* | 465 | 0 | Y | Y | Y |
| Fabaceae | *Vicia sativa* | 208 | 0 | Y | Y | Y |
| Gentianaceae | *Centaurium* spp. | 2 | 0 | | Y | Y |
| Gentianaceae | *Sabatia campestris* | 18 | 0 | | | Y |
| Geraniaceae | *Erodium cicutarium* | 1 | 2 | Y | | |
| Geraniaceae | *Geranium carolinianum* | 10 | 0 | Y | Y | |
| Geraniaceae | *Geranium texanum* | 72 | 0 | Y | Y | Y |
| Heliotropiaceae | *Heliotropium tenellum* | 162 | 1 | Y | Y | Y |
| Hypoxidaceae | *Hypoxis hirsuta* | 0 | 0 | Y | | |
| Iridaceae | *Nemastylis geminiflora* | 0 | 1 | Y | | Y |
| Iridaceae | *Sisyrinchium* spp. | 302 | 11 | Y | Y | Y |
| Krameriaceae | *Krameria lanceolata* | 144 | 0 | Y | Y | Y |
| Lamiaceae | *Hedeoma drummondii* | 16 | 0 | Y | | Y |
| Lamiaceae | *Hedeoma hispida* | 3 | 0 | | | Y |
| Lamiaceae | *Lamium amplexicaule* | 2 | 0 | Y | | |
| Lamiaceae | *Monarda citriodora* | 768 | 103 | Y | Y | Y |

| Family | Taxon | Abundance in quadrats | # observed insect visits | Present at continuous sites? | Present at low frequency rotation sites? | Present at high frequency rotation sites? |
|---|---|---|---|---|---|---|
| Lamiaceae | *Salvia azurea* | 2 | 0 | | | Y |
| Lamiaceae | *Salvia engelmannii* | 0 | 0 | | Y | |
| Lamiaceae | *Scutellaria drummondii* | 191 | 6 | Y | Y | Y |
| Linaceae | *Linum pratense* | 72 | 3 | Y | Y | Y |
| Linaceae | *Linum rigidum* | 30 | 10 | Y | Y | Y |
| Linaceae | *Linum sulcatum* | 7 | 0 | | Y | Y |
| Lythraceae | *Lythrum alatum* | 17 | 0 | | | Y |
| Lythraceae | *Lythrum californicum* | 18 | 1 | | | Y |
| Malvaceae | *Callirhoe alcaeoides* | 0 | 1 | | Y | Y |
| Malvaceae | *Callirhoe involucrata* | 1 | 0 | Y | | |
| Malvaceae | *Callirhoe pedata* | 2 | 1 | | | Y |
| Malvaceae | *Sida abutilifolia* | 7 | 0 | Y | Y | Y |
| Montiaceae | *Claytonia virginica* | 3 | 1 | | | Y |
| Nyctaginaceae | *Mirabilis albida/linearis* | 0 | 0 | | | Y |
| Onagraceae | *Oenothera berlandieri* | 36 | 9 | Y | Y | Y |
| Onagraceae | *Oenothera glaucifolia* | 12 | 9 | | | Y |
| Onagraceae | *Oenothera laciniata* | 1 | 0 | Y | | |
| Onagraceae | *Oenothera speciosa* | 0 | 7 | Y | Y | Y |
| Onagraceae | *Oenothera suffulta* | 249 | 0 | Y | Y | Y |
| Orobanchaceae | *Agalinis heterophylla* | 0 | 1 | Y | | Y |
| Orobanchaceae | *Castilleja indivisa* | 66 | 0 | | | Y |
| Orobanchaceae | *Castilleja purpurea* | 3 | 0 | | | Y |
| Oxalidaceae | *Oxalis dillenii* | 25 | 0 | Y | Y | Y |
| Phyllanthaceae | *Phyllanthus polygonoides* | 597 | 0 | Y | Y | Y |
| Phytolaccaceae | *Rivina humilis* | 0 | 0 | Y | | |

| Family | Taxon | Abundance in quadrats | # observed insect visits | Present at continuous sites? | Present at low frequency rotation sites? | Present at high frequency rotation sites? |
|---|---|---|---|---|---|---|
| Plantaginaceae | *Nuttallanthus texanus* | 4 | 0 | Y | | |
| Plantaginaceae | *Penstemon cobaea* | 0 | 0 | | | Y |
| Plantaginaceae | *Plantago virginica* | 320 | 1 | Y | Y | Y |
| Plantaginaceae | *Veronica arvensis* | 253 | 0 | Y | Y | Y |
| Plantaginaceae | *Veronica peregrina* | 0 | 0 | | Y | |
| Polemoniaceae | *Phlox pilosa* | 105 | 0 | | Y | Y |
| Polygalaceae | *Polygala alba* | 15 | 0 | | | Y |
| Polygonaceae | *Eriogonum longifolium* | 50 | 2 | | | Y |
| Polygonaceae | *Rumex pulcher* | 0 | 0 | Y | | |
| Ranunculaceae | *Anemone berlandieri* | 150 | 24 | Y | Y | Y |
| Ranunculaceae | *Delphinium carolinianum* ssp. *virescens* | 0 | 0 | | Y | Y |
| Ranunculaceae | *Ranunculus* spp. | 0 | 0 | Y | | |
| Rhamnaceae | *Ceanothus herbaceus* | 4 | 1 | | Y | Y |
| Rosaceae | *Rosa foliolosa* | 7 | 9 | | | Y |
| Rubiaceae | *Cruciata pedemontana* | 29 | 0 | | | Y |
| Rubiaceae | *Galium aparine* | 299 | 0 | Y | Y | Y |
| Rubiaceae | *Galium virgatum* | 10738 | 0 | Y | Y | Y |
| Rubiaceae | *Houstonia pusilla* | 1 | 0 | Y | | |
| Rubiaceae | *Sherardia arvensis* | 140 | 0 | Y | | Y |
| Rubiaceae | *Stenaria nigricans* | 6473 | 29 | Y | | Y |
| Solanaceae | *Physalis* spp. | 0 | 0 | Y | | |
| Solanaceae | *Solanum dimidiatum* | 15 | 17 | Y | Y | Y |
| Valerianaceae | *Valerianella amarella/radiata* | 10338 | 17 | Y | Y | Y |
| Verbenaceae | *Glandularia bipinnatifida* | 56 | 17 | Y | Y | Y |
| Verbenaceae | *Verbena halei* | 2 | 0 | Y | | Y |

| Family | Taxon | Abundance in quadrats | # observed insect visits | Present at continuous sites? | Present at low frequency rotation sites? | Present at high frequency rotation sites? |
|---|---|---|---|---|---|---|
| Violaceae | *Hybanthus verticillatus* | 2 | 0 | Y | | |
| Violaceae | *Viola bicolor* | 5 | 0 | Y | | Y |

**Table S2:** Taxonomic resources used to identify insects. Taxonomic experts who provided or verified identifications included John Ascher (bees), Val Bugh (Diptera), Michael Caballero (Coleoptera), Bill Dempwolf (Lepidoptera), Rick Fleischer (Lepidoptera), Jason Hansen (Coleoptera), James McDermott (Lepidoptera), Ed G. Riley (Coleoptera), and Ashley J. Schmitz (Coleoptera, Hymenoptera). Morphospecies names are unique taxa that have not yet been formally described (e.g., "sp. TX-03"). Names follow those used by regional taxonomic experts.

| **Order** | **Family** | **Genus, species, species group** | **Identification resources** |
|---|---|---|---|
| Coleoptera | Anthribidae | *Trigonorhinus limbatus-tomentosus* | (Valentine, 1998) |
| | Attelabidae | *Haplorhynchites pseudomexicanus* | (Hamilton, 1974) |
| | Bupresidae | *Acmaeodera mixta*, *Acmaeodera ornatoides*, *Acmaeodera tubulus* | (Hansen, 2023) |
| | Cantharidae | *Chauliognathus scutellaris* | (Fender, 1964) |
| | Carabidae | *Lebia viridis* | (Horn, 1872) |
| | Cerambycidae | *Batyle suturalis* | (Evans, 2014) |
| | Cerambycidae | *Typocerus octonotatus* | (Yanega, 1996) |
| | Chrysomelidae | *Algarobius bottimeri* | (Kingsolver, 2004a, 2004b) |
| | Chrysomelidae | *Brachypnoea lecontei* | (E.G. Riley, pers. comm.) |
| | Chrysomelidae | *Chrysolina auripennis* | (Van Dyke, 1937) |
| | Chrysomelidae | *Diabrotica cristata*, *Diabrotica undecimpunctata* | (Krysan, 1986) |
| | Chrysomelidae | *Metrioidea popenoei* | (E.G. Riley, pers. comm.) |
| | Chrysomelidae | *Sennius cruentatus* | (Kingsolver, 2004a, 2004b) |
| | Coccinellidae | *Coccinella septempunctata*, *Diomus terminatus*, *Hippodamia convergens* | (Gordon, 1985) |
| | Curculionidae | *Odontocorynus salebrosus* | (Prena, 2008) |
| | Curculionidae | *Rhyssomatus lineaticollis* | (Blatchley and Leng, 1916) |
| | Curculionidae | *Smicronyx* spp. | (Anderson, 1962) |
| | Dermestidae | *Cryptorhopalum* spp. | (Beal, 1985) |

| | | | |
|---|---|---|---|
| | Meloidae | *Epicauta callosa*, *Epicauta ferruginea*, *Epicauta immaculata*, *Epicauta pensylvanica* | (Dillon, 1952; Pinto and Bologna, 1999; Schmidt, 2008) |
| | Meloidae | *Nemognatha piazata bicolor* | (Dillon, 1952; Pinto and Bologna, 1999; Schmidt, 2008) |
| | Meloidae | *Pyrota perversa* | (Dillon, 1952; Pinto and Bologna, 1999) |
| | Mordellidae | *Mordella atrata*, *Mordellistena cervicalis* | (Liljeblad, 1945) |
| | Phalacridae | *Stilbus apicalis* | (Gimmel, 2011) |
| | Scarabaeidae | *Euphoria kernii* | (Orozco, 2012) |
| | Scarabaeidae | *Trichiotinus piger*, *Trichiotinus texanus* | (Philips et al., 2016; E.G. Riley, pers. comm.) |
| Diptera | Anthomyiidae | Anthomyiidae spp., *Leucophora unilineata* | (Huckett, 1940; McAlpine et al., 1993, 1981) |
| | Bibionidae | *Dilophus orbatus*, *Dilophus* spp. | (Dankowicz, 2023; McAlpine et al., 1981) |
| | Bombyliidae | *Geron* spp., *Villa* spp. | (McAlpine et al., 1981) |
| | Bombyliidae | *Paravilla separata*, *Systoechus solitus*, *Systoechus vulgaris* | (Kits et al., 2008) |
| | Bombyliidae | *Poecilanthrax lucifer*, *Poecilognathus* spp. | (Niu et al., 2025) |
| | Calliphoridae | *Cochliomyia macellaria*, *Lucilia illustris* | (McAlpine et al., 1993) |
| | Chloropoidae | *Apotropina* spp., Chloropidae spp. | (McAlpine et al., 1981) |
| | Drosophilidae | Drosophilidae spp. | (McAlpine et al., 1981) |
| | Hybotidae | Hybotidae spp. | (McAlpine et al., 1981) |
| | Hybotidae | Tachydromiinae spp. | (McAlpine et al., 1981) |
| | Muscidae | *Polietes* spp. | (McAlpine et al., 1993) |
| | Muscidae | *Stomoxys calcitrans* | (McAlpine et al., 1981) |
| | Sarcophagidae | Miltogramminae spp., Sarcophagidae spp., Sarcophaginae spp. | (McAlpine et al., 1993) |
| | Sciaridae | Sciaridae spp. | (McAlpine et al., 1981) |
| | Sepsidae | *Sepsis* spp. | (McAlpine et al., 1993; Melander and Spuler, 1917) |

| | | | |
|---|---|---|---|
| | Syrphidae | *Dioprosopa clavata*, *Eupeodes volucris*, *Paragus haemorrhous* | (McAlpine et al., 1993; Miranda et al., 2013) |
| | Syrphidae | *Eristalis stipator*, *Toxomerus marginatus*, *Toxomerus* spp. | (Dankowicz and Dankowicz, 2021; McAlpine et al., 1993; Miranda et al., 2013) |
| | Stratiomyidae | *Psellidotus hieroglyphicus-johnsoni*, *Psellidotus viridis* | (McAlpine et al., 1981) |
| | Tachinidae | *Archytas apicifer*, *Archytas californiae*, *Gonia* spp., *Peleteria* spp., *Prospherysa* spp., *Spallanzania* spp., *Tachina* spp., Tachinidae spp. | (McAlpine et al., 1993; A.J. Schmidt, pers. comm.) |
| | Tabanidae | *Esenbeckia* spp. | (McAlpine et al., 1981) |
| | Tephritidae | *Neotephritis finalis* | (McAlpine et al., 1993) |
| | Tipulidae | Tipulinae spp. | (McAlpine et al., 1981) |
| Hymenoptera | Anthophila (bees) | | (Michener, 2007; Michener et al., 1994; Mitchell, 1962, 1960) |
| | Andrenidae | *Andrena crawfordi*, *Andrena gardineri*, *Andrena nothoscordi*, *Andrena sitiliae* | (LaBerge, 1967, 1986, 1985) |
| | Andrenidae | *Perdita ignota* | (Timberlake, 1956) |
| | Apidae | *Apis mellifera* | (Mitchell, 1962) |
| | Apidae | *Bombus griseocollis*, *Bombus pensylvanicus* | (Williams et al., 2014) |
| | Apidae | *Ceratina strenua* | (Daly, 1973) |
| | Apidae | *Eucera speciosa* | (Timberlake, 1969) |
| | Apidae | *Melissodes boltoniae-tinctus*, *Melissodes coreopsis*, *Melissodes tepaneca*, *Melissodes* spp. | (LaBerge, 1961, 1957, 1956) |
| | Apidae | *Xylocopa virginica* | (Hurd, 1961) |
| | Argidae | *Schizocerella pilicornis* | (Hartsough et al., 2007) |
| | Braconidae | Braconidae spp. | (Wharton et al., 1997) |
| | Chalcididae | *Conura albifrons* | (Delvare and Bouček, 1992) |
| | Crabronidae | *Bembix americana* | (Parker, 1917) |

| | | | |
|---|---|---|---|
| | Colletidae | *Colletes birkmanni* | (Stephens, 1954; Timberlake, 1951) |
| | Colletidae | *Hylaeus floridanus* | (Snelling, 1970) |
| | Halictidae | *Agapostemon angelicus*, *Agapostemon sericeus* | (Portman et al., 2024; Roberts, 1972) |
| | Halictidae | *Halictus ligatus* | (Roberts, 1973; Timberlake, 1961) |
| | Halictidae | *Lasioglossum connexum*, *Lasioglossum disparile*, *Lasioglossum hudsoniellum*, *Lasioglossum imitatum*, *Lasioglossum semicaeruleum*, *Lasioglossum* sp. TX-24, *Lasioglossum* sp. TX-03, *Lasioglossum* sp. TX-06, *Lasioglossum* sp. TX-16, *Lasioglossum* spp. | (Gardner and Gibbs, 2023, 2020; Gibbs, 2011, 2010; Gibbs et al., 2013) |
| | Ichneumonidae | Campopleginae spp. | (Santos, 2025) |
| | Ichneumonidae | *Compsocryptus texensis* | (Broad, 2011; Townes and Townes, 1962) |
| | Megachilidae | *Coelioxys octodentatus* | (Baker and Baker, 1975) |
| | Megachilidae | *Hoplitis pilosifrons* | (Michener, 1955) |
| | Megachilidae | *Lithurgopsis gibbosa* | (Mitchell, 1938; Snelling, 1983) |
| | Megachilidae | *Megachile amica*, *Megachile brevis*, *Megachile comata*, *Megachile inimica*, *Megachile parallela*, *Megachile policaris* | (Bzdyk, 2012; Gonzalez and Griswold, 2007; Mitchell, 1937, 1933; Sheffield, 2013) |
| | Megachilidae | *Osmia chalybea*, *Osmia subfasciata* | (Griswold and Rightmyer, 2017) |
| | Melittidae | *Hesperapis rodecki* | (Stage, 1966) |
| | Perilampidae | Perilampidae spp. | (Wharton et al., 1997) |
| | Philanthidae | *Cerceris bicornuta* | (Buck et al., 2005) |
| | Pompilidae | *Anoplius aethiops*, *Anoplius depressipes* | (Evans, 1950) |
| | Pompilidae | *Entypus fulvicornis* | (Townes, 1957) |
| | Scoliidae | *Dielis plumipes* | (Grissell, 2013) |
| | Sphecidae | *Isodontia mexicana* | (Brimley, 1938) |
| | Sphecidae | *Prionyx atratus* | (Parker, 1960) |

| | | | |
|---|---|---|---|
| | Thynnidae | *Myzinum quinquecinctum* | (Kimsey, 2009) |
| | Vespidae | *Euodynerus annulatus*, *Stenodynerus ammonia* | (Buck et al., 2008) |
| | Vespidae | *Polistes apachus*, *Polistes bellicosus*, *Polistes carolina*, *Polistes dorsalis*, *Polistes fuscatus*, *Polistes metricus* | (Buck et al., 2012, 2008) |
| Lepidoptera | Erebidae | *Caenurgia* spp. | (J. McDermott, pers. comm.) |
| | Hesperiidae | *Amblyscirtes eos*, *Atalopedes huron*, *Burnsius albescens-communis*, *Erynnis horatius* | (Abbott and Abbott, 2020) |
| | Hesperiidae | *Lerodea eufala*, *Polites otho* | (R. Fleischer, pers. comm.) |
| | Lycaenidae | *Echinargus isola*, *Strymon melinus* | (Abbott and Abbott, 2020) |
| | Nymphalidae | *Agraulis vanilla*, *Asterocampa celtis*, *Danaus plexippus*, *Euptoieta claudia*, *Junonia coenia*, *Vanessa cardui*, *Vanessa virginiensis* | (Abbott and Abbott, 2020) |
| | Papilionidae | *Papilio polyxenes* | (Abbott and Abbott, 2020) |
| | Pieridae | *Abaeis nicippe*, *Colias eurytheme*, *Nathalis iole*, *Phoebis sennae*, *Pontia protodice* | (Abbott and Abbott, 2020) |
| | Psychidae | *Acoloithus falsarius* | (R. Fleischer, pers. comm.) |
| | Scythrididae | Scythrididae spp. | (J. McDermott, pers. comm.) |

**Table S3**: Abundance of each insect taxon we collected, and the management practice(s) under which it was found. Morphospecies names are unique taxa that have not yet been formally described (e.g., "sp. TX-03"). Names follow those used by regional taxonomic experts. We also included several species groups of closely-related species that cannot consistently be distinguished. Richness calculations and network analyses excluded several damaged specimens that could not be identified with sufficient taxonomic resolution to distinguish them as the same or different from other specimens that were identified to a finer resolution (e.g., Coleoptera spp. Diptera spp., Hymenoptera spp., Lepidoptera spp., Sarcophagidae spp., *Lasioglossum* spp., *Toxomerus* spp.).

| **Order** | **Taxon** | **Abundance** | **Present at continuous sites?** | **Present at low frequency rotation sites?** | **Present at high frequency rotation sites?** |
|---|---|---|---|---|---|
| Coleoptera | *Acmaeodera mixta* | 5 | Y | | Y |
| Coleoptera | *Acmaeodera ornatoides* | 7 | | | Y |
| Coleoptera | *Acmaeodera tubulus* | 7 | Y | Y | Y |
| Coleoptera | *Algarobius bottimeri* | 1 | | Y | |
| Coleoptera | *Batyle suturalis* | 2 | Y | | Y |
| Coleoptera | *Brachypnoea lecontei* | 1 | | Y | |
| Coleoptera | *Chauliognathus scutellaris* | 1 | | | Y |
| Coleoptera | *Chrysolina auripennis* | 1 | | | Y |
| Coleoptera | *Coccinella septempunctata* | 4 | Y | Y | Y |
| Coleoptera | Coleoptera spp. | 3 | | | Y |
| Coleoptera | *Cryptorhopalum* spp. | 663 | Y | Y | Y |
| Coleoptera | *Diabrotica cristata* | 9 | | | Y |
| Coleoptera | *Diabrotica undecimpunctata* | 13 | Y | | Y |
| Coleoptera | *Diomus terminatus* | 1 | | Y | |
| Coleoptera | *Epicauta callosa* | 14 | Y | Y | Y |
| Coleoptera | *Epicauta ferruginea* | 4 | Y | | Y |
| Coleoptera | *Epicauta immaculata* | 1 | | Y | |
| Coleoptera | *Epicauta pensylvanica* | 12 | Y | Y | Y |
| Coleoptera | *Euphoria kernii* | 9 | Y | | Y |
| Coleoptera | *Haplorhynchites pseudomexicanus* | 1 | | | Y |
| Coleoptera | *Hippodamia convergens* | 10 | Y | Y | Y |
| Coleoptera | *Lebia viridis* | 1 | | Y | |
| Coleoptera | *Metrioidea popenoei* | 20 | | | Y |

| Order | Taxon | Abundance | Present at continuous sites? | Present at low frequency rotation sites? | Present at high frequency rotation sites? |
|---|---|---|---|---|---|
| Coleoptera | *Mordella atrata* | 1 | Y | | |
| Coleoptera | *Mordellistena cervicalis* | 28 | Y | | Y |
| Coleoptera | *Nemognatha piazata bicolor* | 3 | Y | | Y |
| Coleoptera | *Odontocorynus salebrosus* | 162 | Y | Y | Y |
| Coleoptera | *Pyrota perversa* | 1 | | | Y |
| Coleoptera | *Rhyssomatus lineaticollis* | 1 | | | Y |
| Coleoptera | *Sennius cruentatus* | 2 | Y | | |
| Coleoptera | *Smicronyx* spp. | 5 | | | Y |
| Coleoptera | *Stilbus apicalis* | 1 | | Y | |
| Coleoptera | *Trichiotinus piger* | 1 | | | Y |
| Coleoptera | *Trichiotinus texanus* | 2 | | | Y |
| Coleoptera | *Trigonorhinus limbatus-tomentosus* | 5 | Y | | Y |
| Coleoptera | *Typocerus octonotatus* | 16 | Y | | Y |
| Diptera | Anthomyiidae spp. | 5 | Y | Y | Y |
| Diptera | *Apotropina* spp. | 1 | | Y | |
| Diptera | *Archytas apicifer* | 14 | Y | | Y |
| Diptera | *Archytas californiae* | 10 | | | Y |
| Diptera | Chloropidae spp. | 8 | Y | Y | Y |
| Diptera | *Cochliomyia macellaria* | 1 | | | Y |
| Diptera | *Dilophus orbatus* | 5 | Y | Y | Y |
| Diptera | *Dilophus* spp. | 2 | Y | Y | |
| Diptera | *Dioprosopa clavata* | 1 | | | Y |
| Diptera | Diptera spp. | 6 | Y | Y | Y |
| Diptera | Drosophilidae spp. | 1 | | | Y |
| Diptera | *Eristalis stipator* | 5 | Y | | Y |
| Diptera | *Esenbeckia spp.* | 1 | | | Y |
| Diptera | *Eupeodes volucris* | 4 | Y | Y | |
| Diptera | *Geron* spp. | 20 | Y | Y | Y |
| Diptera | *Gonia* spp. | 1 | | | Y |
| Diptera | Hybotidae spp. | 1 | | | Y |
| Diptera | *Leucophora unilineata* | 1 | | | Y |
| Diptera | *Lucilia illustris* | 25 | Y | Y | Y |
| Diptera | Miltogramminae spp. | 1 | | | Y |

| Order | Taxon | Abundance | Present at continuous sites? | Present at low frequency rotation sites? | Present at high frequency rotation sites? |
|---|---|---|---|---|---|
| Diptera | *Neotephritis finalis* | 1 | | Y | |
| Diptera | *Paragus haemorrhous* | 1 | | | Y |
| Diptera | *Paravilla separata* | 1 | Y | | |
| Diptera | *Peleteria* spp. | 1 | | | Y |
| Diptera | *Poecilanthrax lucifer* | 2 | | | Y |
| Diptera | *Poecilognathus* spp. | 1 | | | Y |
| Diptera | *Polietes* spp. | 1 | | | Y |
| Diptera | *Prospherysa* spp. | 1 | | | Y |
| Diptera | *Psellidotus hieroglyphicus-johnsoni* | 2 | | | Y |
| Diptera | *Psellidotus viridis* | 1 | | | Y |
| Diptera | Sarcophagidae spp. | 1 | | | Y |
| Diptera | Sarcophaginae spp. | 7 | | Y | Y |
| Diptera | Sciaridae spp. | 242 | Y | Y | Y |
| Diptera | *Sepsis* spp. | 3 | Y | Y | Y |
| Diptera | *Spallanzania* spp. | 1 | | | Y |
| Diptera | *Stomoxys calcitrans* | 3 | | | Y |
| Diptera | *Systoechus solitus* | 2 | | Y | |
| Diptera | *Systoechus vulgaris* | 9 | Y | | Y |
| Diptera | *Tachina* spp. | 1 | | | Y |
| Diptera | Tachinidae spp. | 1 | | | Y |
| Diptera | Tachydromiinae spp. | 1 | | Y | |
| Diptera | Tipulinae spp. | 1 | Y | | |
| Diptera | *Toxomerus marginatus* | 54 | Y | Y | Y |
| Diptera | *Toxomerus* spp. | 1 | | | Y |
| Diptera | *Villa* spp. | 2 | | | Y |
| Hymenoptera | *Agapostemon angelicus* | 2 | Y | | Y |
| Hymenoptera | *Agapostemon sericeus* | 1 | | | Y |
| Hymenoptera | *Andrena crawfordi* | 9 | | Y | Y |
| Hymenoptera | *Andrena gardineri* | 1 | | | Y |
| Hymenoptera | *Andrena nothoscordi* | 5 | | | Y |
| Hymenoptera | *Andrena sitiliae* | 2 | | | Y |
| Hymenoptera | *Anoplius aethiops* | 1 | Y | | |
| Hymenoptera | *Anoplius depressipes* | 1 | Y | | |
| Hymenoptera | *Apis mellifera* | 120 | Y | Y | Y |

| Order | Taxon | Abundance | Present at continuous sites? | Present at low frequency rotation sites? | Present at high frequency rotation sites? |
|---|---|---|---|---|---|
| Hymenoptera | *Bembix americana* | 1 | Y | | |
| Hymenoptera | *Bombus griseocollis* | 4 | | Y | Y |
| Hymenoptera | *Bombus pensylvanicus* | 7 | Y | | Y |
| Hymenoptera | Braconidae spp. | 1 | | | Y |
| Hymenoptera | Campopleginae spp. | 1 | Y | | |
| Hymenoptera | *Ceratina strenua* | 3 | | Y | Y |
| Hymenoptera | *Cerceris bicornuta* | 1 | Y | | |
| Hymenoptera | *Coelioxys octodentatus* | 1 | | | Y |
| Hymenoptera | *Colletes birkmanni* | 4 | Y | | Y |
| Hymenoptera | *Compsocryptus texensis* | 1 | Y | | |
| Hymenoptera | *Conura albifrons* | 1 | | | Y |
| Hymenoptera | *Dielis plumipes* | 3 | | | Y |
| Hymenoptera | *Entypus fulvicornis* | 2 | | Y | Y |
| Hymenoptera | *Eucera speciosa* | 1 | Y | | |
| Hymenoptera | *Euodynerus annulatus* | 1 | Y | | |
| Hymenoptera | *Halictus ligatus* | 3 | Y | | Y |
| Hymenoptera | *Hesperapis rodecki* | 1 | | | Y |
| Hymenoptera | *Hoplitis pilosifrons* | 1 | | | Y |
| Hymenoptera | *Hylaeus floridanus* | 1 | Y | | |
| Hymenoptera | Hymenoptera spp. | 2 | | Y | Y |
| Hymenoptera | *Isodontia mexicana* | 3 | | | Y |
| Hymenoptera | *Lasioglossum connexum* | 1 | | Y | |
| Hymenoptera | *Lasioglossum disparile* | 11 | Y | Y | Y |
| Hymenoptera | *Lasioglossum hudsoniellum* | 3 | | Y | Y |
| Hymenoptera | *Lasioglossum imitatum* | 1 | Y | | |
| Hymenoptera | *Lasioglossum semicaeruleum* | 6 | Y | Y | Y |
| Hymenoptera | *Lasioglossum* sp. TX-16 | 3 | | Y | Y |
| Hymenoptera | *Lasioglossum* sp. TX-24 | 3 | | | Y |
| Hymenoptera | *Lasioglossum* sp. TX-3 | 6 | | Y | Y |
| Hymenoptera | *Lasioglossum* sp. TX-06 | 2 | | Y | Y |
| Hymenoptera | *Lasioglossum* spp. | 1 | | | Y |
| Hymenoptera | *Lithurgopsis gibbosa* | 2 | | | Y |
| Hymenoptera | *Megachile amica* | 1 | Y | | |

| Order | Taxon | Abundance | Present at continuous sites? | Present at low frequency rotation sites? | Present at high frequency rotation sites? |
|---|---|---|---|---|---|
| Hymenoptera | *Megachile brevis* | 7 | Y | Y | Y |
| Hymenoptera | *Megachile comata* | 2 | | | Y |
| Hymenoptera | *Megachile inimica* | 3 | | | Y |
| Hymenoptera | *Megachile parallela* | 1 | | | Y |
| Hymenoptera | *Megachile policaris* | 1 | | | Y |
| Hymenoptera | *Melissodes boltoniae-tinctus* | 7 | | | Y |
| Hymenoptera | *Melissodes coreopsis* | 10 | Y | Y | Y |
| Hymenoptera | *Melissodes* spp. | 3 | Y | | Y |
| Hymenoptera | *Melissodes tepaneca* | 1 | Y | | |
| Hymenoptera | *Myzinum quinquecinctum* | 2 | | | Y |
| Hymenoptera | *Osmia chalybea* | 1 | | | Y |
| Hymenoptera | *Osmia subfasciata* | 1 | Y | | |
| Hymenoptera | *Perdita ignota* | 1 | Y | | |
| Hymenoptera | Perilampidae spp. | 1 | | | Y |
| Hymenoptera | *Polistes apachus* | 3 | | Y | Y |
| Hymenoptera | *Polistes bellicosus* | 2 | Y | | Y |
| Hymenoptera | *Polistes carolina* | 1 | | | Y |
| Hymenoptera | *Polistes dorsalis* | 2 | Y | | Y |
| Hymenoptera | *Polistes fuscatus* | 9 | | Y | Y |
| Hymenoptera | *Polistes metricus* | 1 | | | Y |
| Hymenoptera | *Prionyx atratus* | 2 | Y | | Y |
| Hymenoptera | *Schizocerella pilicornis* | 1 | | | Y |
| Hymenoptera | *Stenodynerus ammonia* | 1 | | | Y |
| Hymenoptera | *Xylocopa virginica* | 19 | Y | Y | Y |
| Lepidoptera | *Abaeis nicippe* | 3 | Y | | Y |
| Lepidoptera | *Acoloithus falsarius* | 11 | | | Y |
| Lepidoptera | *Agraulis vanillae* | 1 | | | Y |
| Lepidoptera | *Amblyscirtes eos* | 1 | | | Y |
| Lepidoptera | *Asterocampa celtis* | 1 | Y | | |
| Lepidoptera | *Atalopedes huron* | 23 | Y | Y | Y |
| Lepidoptera | *Burnsius albescens-communis* | 2 | | | Y |
| Lepidoptera | *Caenurgia* spp. | 1 | Y | | |
| Lepidoptera | *Colias eurytheme* | 26 | Y | Y | Y |

| Order | Taxon | Abundance | Present at continuous sites? | Present at low frequency rotation sites? | Present at high frequency rotation sites? |
|---|---|---|---|---|---|
| Lepidoptera | *Danaus plexippus* | 22 | Y | | Y |
| Lepidoptera | *Echinargus isola* | 1 | | Y | |
| Lepidoptera | *Erynnis horatius* | 2 | | | Y |
| Lepidoptera | *Euptoieta claudia* | 16 | Y | Y | Y |
| Lepidoptera | *Junonia coenia* | 17 | Y | Y | Y |
| Lepidoptera | Lepidoptera spp. | 1 | | Y | |
| Lepidoptera | *Lerodea eufala* | 1 | Y | | |
| Lepidoptera | *Nathalis iole* | 21 | Y | | Y |
| Lepidoptera | *Papilio polyxenes* | 1 | Y | | |
| Lepidoptera | *Phoebis sennae* | 1 | | Y | |
| Lepidoptera | *Polites otho* | 1 | Y | | |
| Lepidoptera | *Pontia protodice* | 3 | Y | Y | Y |
| Lepidoptera | Scythrididae spp. | 6 | | | Y |
| Lepidoptera | *Strymon melinus* | 16 | Y | Y | Y |
| Lepidoptera | *Vanessa cardui* | 1 | | | Y |
| Lepidoptera | *Vanessa virginiensis* | 1 | | | Y |

19

**Table S4:** Structural equation models assessing impacts of soil texture, management, and flower abundance on pollinator abundance. SEMs asked how management strategy (continuous stocking, low-frequency rotation, high-frequency rotation) and soil texture (as sand content) directly and indirectly via flower abundance impacted abundance of (1) all pollinator taxa combined, (2) all pollinator taxa except for the highly abundant *Cryptorhopalum* dermestid beetles, (3) Coleoptera, (4) Coleoptera without *Cryptorhopalum*, (5) Diptera, (6) Hymenoptera, and (7) Lepidoptera. The flower abundance portion was invariant across the SEMs. Terms significant at α=0.05 are bolded.

| **Response variable** | **Predictor** | **Standardized coefficient** | **Coefficient standard error** | **Degrees of freedom** | **$X^2$** | **p-value** | **Continuous stocking coefficient (lower & upper confidence limits)** | **Low frequency rotation coefficient (lower & upper confidence limits)** | **High frequency rotation coefficient (lower & upper confidence limits)** |
|---|---|---|---|---|---|---|---|---|---|
| Flower abundance | Soil sand content | 0.02 | 0.02 | 15 | 0.97 | 0.33 | | | |
| | Management strategy | | | 2 | 9.56 | **0.002** | 7.24 (6.42, 8.06) | 7.29 (6.42, 8.16) | 9.11 (8.43, 9.80) |
| Pollinator abundance | Soil sand content | -0.04 | 0.01 | 14 | -4.50 | **<0.0001** | | | |
| | Flower abundance | 0.0001 | 0 | 14 | 2.18 | **0.03** | | | |
| | Management strategy | | | 2 | 0.12 | 0.89 | 4.62 (4.27, 4.97) | 4.60 (4.25, 4.95) | 4.47 (4.13, 4.81) |
| Pollinator abundance without *Cryptorhopalum* | Soil sand content | -0.02 | 0.01 | 14 | -2.04 | **0.04** | | | |
| | Flower abundance | 0 | 0 | 14 | 1.37 | 0.17 | | | |

| **Response variable** | **Predictor** | **Standardized coefficient** | **Coefficient standard error** | **Degrees of freedom** | **$X^2$** | **p-value** | **Continuous stocking coefficient (lower & upper confidence limits)** | **Low frequency rotation coefficient (lower & upper confidence limits)** | **High frequency rotation coefficient (lower & upper confidence limits)** |
|---|---|---|---|---|---|---|---|---|---|
| | Management strategy | | | 2 | 0.70 | 0.51 | 3.88 (3.48, 4.29) | 4.02 (3.61, 4.22) | 4.37 (3.99, 4.76) |
| Coleoptera abundance | Soil sand content | -0.07 | 0.01 | 14 | -5.15 | **<0.0001** | | | |
| | Flower abundance | 0.0001 | 0 | 14 | 2.30 | **0.02** | | | |
| | Management strategy | | | 2 | 2.25 | 0.14 | 4.26 (3.75, 4.78) | 4.01 (3.49, 4.53) | **0.27** (-0.25, 0.77) |
| Coleoptera abundance without *Cryptorhopalum* | Soil sand content | -0.06 | 0.02 | 14 | -3.22 | **0.001** | | | |
| | Flower abundance | 0.0001 | 0 | 14 | 2.77 | **0.01** | | | |
| | Management strategy | | | 2 | 0.56 | 0.58 | 3.01 (2.46, 3.71) | 2.84 (2.21, 3.48) | 2.46 (1.85, 3.08) |
| Diptera abundance | Soil sand content | -0.01 | 0.02 | 14 | -0.39 | 0.70 | | | |
| | Flower abundance | 0 | 0.0001 | 14 | -0.17 | 0.87 | | | |
| | Management strategy | | | 2 | 1.35 | 0.29 | 2.25 (1.49, 3.01) | 3.06 (2.32, 3.80) | 3.59 (2.89, 4.30) |
| Hymenoptera abundance | Soil sand content | -0.01 | 0.01 | 14 | -0.72 | 0.47 | | | |

| Response variable | Predictor | Standardized coefficient | Coefficient standard error | Degrees of freedom | $X^2$ | p-value | Continuous stocking coefficient (lower & upper confidence limits) | Low frequency rotation coefficient (lower & upper confidence limits) | High frequency rotation coefficient (lower & upper confidence limits) |
|---|---|---|---|---|---|---|---|---|---|
| | Flower abundance | 0 | 0 | 14 | -0.26 | 0.179 | | | |
| | Management strategy | | | 2 | 1.97 | 0.18 | 2.54 (1.94, 3.15) | 2.04 (1.40, 2.67) | 3.19 (2.62, 3.75) |
| Lepidoptera abundance | Soil sand content | -0.01 | 0.01 | 14 | -0.68 | 0.49 | | | |
| | Flower abundance | 0.0001 | 0 | 14 | 3.32 | **0.0009** | | | |
| | Management strategy | | | 2 | 1.10 | 0.91 | 1.91 (1.34, 2.47) | 1.95 (1.39, 2.51) | 2.14 (1.64, 2.63) |

**Table S5:** Multivariate analysis of abundance to determine impacts of soil texture and management strategy on flower community composition. Terms significant at $\alpha=0.05$ are bolded.

| **Model term** | $X^2$ | **Degrees of freedom** | **Residual degrees of freedom** | **p-value** |
|---|---|---|---|---|
| Intercept | | | 18 | |
| Management | 343.2 | 2 | 16 | **0.001** |
| Soil sand content | 300.3 | 1 | 15 | **0.001** |

**Table S6:** Univariate likelihood ratio tests of impacts of soil texture and management strategy on flower species' abundance, from the community composition analysis. Terms significant at $\alpha$=0.05 are bolded.

| **Taxon** | **Model term** | **$X^2$** | **p-value** |
|---|---|---|---|
| *Allium* spp. | Management | 4.60 | 1 |
| | Soil sand content | 0.18 | 1 |
| *Ammoselinum butleri* | Management | 2.70 | 1 |
| | Soil sand content | 0.59 | 1 |
| *Amsonia ciliata* | Management | 4.23 | 1 |
| | Soil sand content | 0.51 | 1 |
| *Anemone berlandieri* | Management | 1.34 | 1 |
| | Soil sand content | 0.53 | 1 |
| *Arenaria serpyllifolia* | Management | 2.23 | 1 |
| | Soil sand content | 3.62 | 0.99 |
| *Chaerophyllum tainturieri* | Management | 1.09 | 1 |
| | Soil sand content | 0.08 | 1 |
| *Coreopsis lanceolata* | Management | 2.51 | 1 |
| | Soil sand content | 0.78 | 1 |
| *Euphorbia longicruris* | Management | 2.15 | 1 |
| | Soil sand content | 3.85 | 0.99 |
| *Galium virgatum* | Management | 4.81 | 1 |
| | Soil sand content | 0.06 | 1 |
| *Lepidium virginicum* | Management | 5.83 | 0.97 |
| | Soil sand content | 10.81 | **0.04** |
| *Linum pratense* | Management | 2.11 | 1 |
| | Soil sand content | 0.30 | 1 |
| *Linum rigidum* | Management | 0.50 | 0.93 |
| | Soil sand content | 0.44 | 1 |
| *Medicago* spp. | Management | 5.27 | 1 |
| | Soil sand content | 0.76 | 1 |
| *Monarda citriodora* | Management | 3.35 | 1 |
| | Soil sand content | 2.08 | 1 |
| *Nothoscordum bivalve* | Management | 0.20 | 1 |
| | Soil sand content | 0.28 | 1 |
| *Phyllanthus polygonoides* | Management | 4.35 | 1 |
| | Soil sand content | 0.80 | 1 |
| *Plantago virginica* | Management | 7.94 | 0.74 |
| | Soil sand content | 1.71 | 1 |
| *Stenaria nigricans* | Management | 0.25 | 1 |
| | Soil sand content | 1.46 | 1 |

| Taxon | Model term | $X^2$ | p-value |
|---|---|---|---|
| *Tragia* spp. | Management | 2.56 | 1 |
| | Soil sand content | 0 | 1 |
| *Valerianella amerella radiata* | Management | 6.02 | 0.97 |
| | Soil sand content | 0.07 | 1 |
| *Veronica arvensis* | Management | 6.50 | 1 |
| | Soil sand content | 0.02 | 0.95 |
| *Vicia sativa* | Management | 3.57 | 1 |
| | Soil sand content | 4.61 | 0.92 |
| *Amphiachyris/Gutierrezia* spp. | Management | 0.47 | 1 |
| | Soil sand content | 4.34 | 0.95 |
| *Asclepias viridis* | Management | 4.73 | 1 |
| | Soil sand content | 0.79 | 1 |
| *Astragalus nuttallianaus* | Management | 5.27 | 1 |
| | Soil sand content | 0.57 | 1 |
| *Bifora americana* | Management | 2.44 | 1 |
| | Soil sand content | 0.41 | 1 |
| *Callirhoe pedata* | Management | 2.94 | 1 |
| | Soil sand content | 0.08 | 1 |
| *Castilleja purpurea* | Management | 2.94 | 1 |
| | Soil sand content | 0.12 | 1 |
| *Croton monothogynus* | Management | 6.17 | 0.97 |
| | Soil sand content | 0.05 | 1 |
| *Cuscuta* spp. | Management | 3.89 | 0.934 |
| | Soil sand content | 0.26 | 1 |
| *Dalea aurea* | Management | 2.94 | 1 |
| | Soil sand content | 0.08 | 1 |
| *Diaperia verna* | Management | 6.26 | 0.97 |
| | Soil sand content | 0.10 | 1 |
| *Dyschoriste linearis* | Management | 2.28 | 1 |
| | Soil sand content | 7.45 | 0.31 |
| *Engelmannia peristenia* | Management | 2.69 | 1 |
| | Soil sand content | 6.03 | 0.64 |
| *Eriogonum longifolium* | Management | 2.19 | 1 |
| | Soil sand content | 0.01 | 1 |
| *Euphorbia spathulate* | Management | 1.50 | 1 |
| | Soil sand content | 2.05 | 1 |
| *Evolvulus nuttallianus* | Management | 1.63 | 1 |
| | Soil sand content | 0.24 | 1 |
| *Gaillardia pulchella* | Management | 1.40 | 1 |

| Taxon | Model term | $X^2$ | p-value |
|---|---|---|---|
| | Soil sand content | 0.02 | 1 |
| *Glandularia bipinnatifida* | Management | 1.38 | 1 |
| | Soil sand content | 0.21 | 1 |
| *Hedeoma drummondii* | Management | 2.68 | 1 |
| | Soil sand content | 6.35 | 0.56 |
| *Heliotropium tenellum* | Management | 3.46 | 1 |
| | Soil sand content | 0.01 | 1 |
| *Lathyrus hirsutus* | Management | 2.82 | 1 |
| | Soil sand content | 0.83 | 1 |
| *Lindheimera texana* | Management | 4.22 | 1 |
| | Soil sand content | 0.008 | 1 |
| *Oenothera berlandieri* | Management | 4.81 | 1 |
| | Soil sand content | 2.40 | 1 |
| *Oenothera suffulta* | Management | 0.24 | 1 |
| | Soil sand content | 0.14 | 1 |
| *Paranychia virginica* | Management | 2.80 | 1 |
| | Soil sand content | 5.30 | 0.82 |
| *Physaria gracilis* | Management | 1.76 | 1 |
| | Soil sand content | 8.32 | 0.18 |
| *Plectocephalus americanus* | Management | 2.68 | 1 |
| | Soil sand content | 0.50 | 1 |
| *Polygala alba* | Management | 2.64 | 1 |
| | Soil sand content | 2.98 | 1 |
| *Polytaenia texana* | Management | 3.03 | 1 |
| | Soil sand content | 0.92 | 1 |
| *Scutellaria drummondii* | Management | 6.07 | 0.97 |
| | Soil sand content | 0.76 | 1 |
| *Sida abutilifolia* | Management | 3.30 | 1 |
| | Soil sand content | 0.25 | 1 |
| *Tetraneuris linearifolia* | Management | 1.54 | 1 |
| | Soil sand content | 0.01 | 1 |
| *Thelesperma filifolium* | Management | 2.14 | 1 |
| | Soil sand content | 0.09 | 1 |
| *Vicia ludoviciana* | Management | 2.89 | 1 |
| | Soil sand content | 4.23 | 0.96 |
| *Matelea biflora* | Management | 1.93 | 1 |
| | Soil sand content | 1.23 | 1 |
| *Asclepias asperula* | Management | 2.88 | 1 |
| | Soil sand content | 0.01 | 1 |

| **Taxon** | **Model term** | **$X^2$** | **p-value** |
|---|---|---|---|
| *Ammoselinum popei* | Management | 6.39 | 0.95 |
| | Soil sand content | 0.18 | 1 |
| *Oxalis dillenii* | Management | 6.83 | 0.92 |
| | Soil sand content | 0.05 | 1 |
| *Sisyrinchium* spp. | Management | 1.84 | 1 |
| | Soil sand content | 6.90 | 0.41 |
| *Torilis nodosa* | Management | 6.00 | 0.97 |
| | Soil sand content | 0.99 | 1 |
| *Achillea millefolium* | Management | 4.00 | 1 |
| | Soil sand content | 2.39 | 1 |
| *Claytonia virginica* | Management | 1.86 | 1 |
| | Soil sand content | 7.39 | 0.32 |
| *Erigeron* spp. | Management | 7.22 | 0.85 |
| | Soil sand content | 0.09 | 1 |
| *Krameria lanceolata* | Management | 7.78 | 0.76 |
| | Soil sand content | 0.24 | 1 |
| *Sherardia arvensis* | Management | 3.02 | 1 |
| | Soil sand content | 0 | 1 |
| *Viola bicolor* | Management | 1.85 | 1 |
| | Soil sand content | 8.35 | 0.18 |
| *Lactuca* spp. | Management | 1.78 | 1 |
| | Soil sand content | 2.19 | 1 |
| *Eryngium leavenworthii* | Management | 2.77 | 1 |
| | Soil sand content | 0.36 | 1 |
| *Euphorbia bicolor* | Management | 3.72 | 1 |
| | Soil sand content | 0.01 | 1 |
| *Geranium texanum* | Management | 0.39 | 1 |
| | Soil sand content | 3.03 | 1 |
| *Mimosa* spp. | Management | 3.88 | 1 |
| | Soil sand content | 0.23 | 1 |
| *Acacia angustissima* | Management | 2.41 | 1 |
| | Soil sand content | 2.24 | 1 |
| *Brickellia eupatorioides* | Management | 2.14 | 1 |
| | Soil sand content | 1.04 | 1 |
| *Dalea compacta* | Management | 1.38 | 1 |
| | Soil sand content | 0.34 | 1 |
| *Lythrum alatum* | Management | 2.07 | 1 |
| | Soil sand content | 1.12 | 1 |
| *Oenothera glaucifolia* | Management | 2.16 | 1 |
| | Soil sand content | 3.13 | 1 |

| **Taxon** | **Model term** | $X^2$ | **p-value** |
|---|---|---|---|
| *Phlox pilosa* | Management | 2.68 | 1 |
| | Soil sand content | 2.93 | 1 |
| *Pyrrhopappus pauciflorus* | Management | 5.04 | 1 |
| | Soil sand content | 1.66 | 1 |
| *Solanum dimidiatum* | Management | 2.07 | 1 |
| | Soil sand content | 0.97 | 1 |
| *Torilis arvensis* | Management | 2.20 | 1 |
| | Soil sand content | 1.63 | 1 |
| *Ceanothus herbaceous* | Management | 2.51 | 1 |
| | Soil sand content | 0.84 | 1 |
| *Lythrum californicum* | Management | 2.51 | 1 |
| | Soil sand content | 0.91 | 1 |
| *Symphotrichum subulatum* var. *ligulatum* | Management | 2.51 | 1 |
| | Soil sand content | 0.92 | 1 |
| *Castilleja indivisa* | Management | 3.75 | 1 |
| | Soil sand content | 3.55 | 1 |
| *Diaperia prolifera* | Management | 1.86 | 1 |
| | Soil sand content | 14.08 | **0.005** |
| *Euphorbia missurica* | Management | 1.69 | 1 |
| | Soil sand content | 1.96 | 1 |
| *Hedeoma hispida* | Management | 3.91 | 1 |
| | Soil sand content | 6.78 | 0.44 |
| *Liatris punctata* | Management | 1.85 | 1 |
| | Soil sand content | 10.98 | **0.03** |
| *Salvia azurea* | Management | 1.86 | 1 |
| | Soil sand content | 6.51 | 0.53 |
| *Stillingia texana* | Management | 1.86 | 1 |
| | Soil sand content | 11.91 | **0.03** |
| *Symphotrichum ericoides* | Management | 6.09 | 0.97 |
| | Soil sand content | 1.08 | 1 |
| *Centaurium* spp. | Management | 1.63 | 1 |
| | Soil sand content | 0.44 | 1 |
| *Croton capitatus* | Management | 2.36 | 1 |
| | Soil sand content | 4.14 | 0.98 |
| *Cruciata pedemontana* | Management | 1.85 | 1 |
| | Soil sand content | 1.71 | 1 |
| *Linum sulcatum* | Management | 1.72 | 1 |
| | Soil sand content | 1.45 | 1 |
| *Sabatia campestris* | Management | 1.85 | 1 |

| **Taxon** | **Model term** | **$X^2$** | **p-value** |
|---|---|---|---|
| | Soil sand content | 1.68 | 1 |
| *Capsella bursa pastoris* | Management | 2.51 | 1 |
| | Soil sand content | 5.86 | 0.69 |
| *Croton glandulosus* | Management | 2.51 | 1 |
| | Soil sand content | 8.29 | 0.18 |
| *Galium aparine* | Management | 1.73 | 1 |
| | Soil sand content | 0.58 | 1 |
| *Geranium carolinianum* | Management | 2.50 | 1 |
| | Soil sand content | 8.86 | 0.14 |
| *Hedypnois cretica* | Management | 2.51 | 1 |
| | Soil sand content | 5.86 | 0.69 |
| *Lamium amplexicaule* | Management | 2.51 | 1 |
| | Soil sand content | 5.86 | 0.69 |
| *Nuttallanthus texanus* | Management | 2.51 | 1 |
| | Soil sand content | 7.32 | 0.34 |
| *Sisymbrium officinale* | Management | 2.51 | 1 |
| | Soil sand content | 7.32 | 0.34 |
| *Stellaria media* | Management | 2.51 | 1 |
| | Soil sand content | 6.76 | 0.46 |
| *Verbena halei* | Management | 2.51 | 1 |
| | Soil sand content | 5.86 | 0.69 |
| *Vicia hirsuta* | Management | 2.51 | 1 |
| | Soil sand content | 13.41 | **0.005** |
| *Packera tampicana* | Management | 5.16 | 1 |
| | Soil sand content | 3.34 | 1 |
| *Packera plattensis* | Management | 2.93 | 1 |
| | Soil sand content | 8.81 | 0.14 |
| *Apocynum cannabinum* | Management | 1.86 | 1 |
| | Soil sand content | 2.64 | 1 |
| *Rudbeckia amplexicaulis* | Management | 1.86 | 1 |
| | Soil sand content | 2.64 | 1 |
| *Croton lindheimeri* | Management | 1.85 | 1 |
| | Soil sand content | 0.52 | 1 |
| *Rosa foliosa* | Management | 1.85 | 1 |
| | Soil sand content | 0.54 | 1 |
| *Hybanthus verticillatus* | Management | 2.51 | 1 |
| | Soil sand content | 0.13 | 1 |

**Table S7:** Structural equation models assessing impacts of soil texture, management, and flower abundance on pollinator richness. SEMs asked how management strategy (continuous stocking, low-frequency rotation, high-frequency rotation) and soil texture (as sand content) directly and indirectly via flower abundance impacted richness of (1) all pollinator taxa combined, (2) all pollinator taxa except for the highly abundant *Cryptorhopalum* dermestid beetles, (3) Coleoptera, (4) Coleoptera without *Cryptorhopalum*, (5) Diptera, (6) Hymenoptera, and (7) Lepidoptera. The flower abundance portion was invariant across the SEMs. Pollinator richness calculations excluded several damaged specimens that could not be identified with sufficient taxonomic resolution to distinguish them as the same or different from other specimens that were identified to a finer resolution (e.g., Coleoptera spp. Diptera spp., Hymenoptera spp., Lepidoptera spp., Sarcophagidae spp., *Lasioglossum* spp., *Toxomerus* spp.). Richness was estimated using the iNEXT R package. Terms significant at $\alpha$=0.05 are bolded.

| **Response variable** | **Predictor** | **Standardized coefficient** | **Coefficient standard error** | **Degrees of freedom** | **$X^2$** | **p-value** | **Continuous stocking coefficient (lower & upper confidence limits)** | **Low frequency rotation coefficient (lower & upper confidence limits)** | **High frequency rotation coefficient (lower & upper confidence limits)** |
|---|---|---|---|---|---|---|---|---|---|
| Flower abundance | Soil sand content | 0.02 | 0.02 | 15 | 0.97 | 0.33 | | | |
| | Management strategy | | | 2 | 9.56 | **0.002** | 7.24 (6.42, 8.06) | 7.29 (6.42, 8.16) | 9.11 (8.43, 9.80) |
| Pollinator richness | Soil sand content | 0.12 | 0.60 | 14 | 0.19 | 0.86 | | | |
| | Flower abundance | -0.0002 | 0.002 | 14 | -0.11 | 0.92 | | | |
| | Management strategy | | | 2 | 2.99 | 0.08 | 70.1 (42.92, 97.37) | 33. 8 (6.53, 61.13) | 77.1 (50.80, 103.35) |
| Pollinator richness without *Cryptorhopalum* | Soil sand content | -0.08 | 0.57 | 14 | -0.13 | 0.90 | | | |

| **Response variable** | **Predictor** | **Standardized coefficient** | **Coefficient standard error** | **Degrees of freedom** | **$X^2$** | **p-value** | **Continuous stocking coefficient (lower & upper confidence limits)** | **Low frequency rotation coefficient (lower & upper confidence limits)** | **High frequency rotation coefficient (lower & upper confidence limits)** |
|---|---|---|---|---|---|---|---|---|---|
| | Flower abundance | 0 | 0.002 | 14 | 0.02 | 0.99 | | | |
| | Management strategy | | | 2 | 3.08 | 0.02 | 68.25 (42.29, 94.21) | 32.93 (6.89, 58.96) | 74.67 (49.62, 99.73) |
| Coleoptera richness | Soil sand content | -0.09 | 0.08 | 14 | -1.20 | 0.25 | | | |
| | Flower abundance | 0.0005 | 0.0002 | 14 | 2.04 | 0.06 | | | |
| | Management strategy | | | 2 | 0.63 | 0.54 | 10.16 (6.59, 13.73) | 7.76 (4.18, 11.35) | 7.59 (4.14, 11.03) |
| Coleoptera richness without *Cryptorhopalum* | Soil sand content | -0.12 | 0.08 | 14 | -1.46 | 0.17 | | | |
| | Flower abundance | 0.0005 | 0.0002 | 14 | 2.22 | **0.04** | | | |
| | Management strategy | | | 2 | 0.64 | 0.54 | 9.02 (54.45, 12.58) | 6.71 (3.14, 10.28) | 6.29 (2.86, 9.73) |
| Diptera richness | Soil sand content | 0.06 | 0.19 | 14 | 0.32 | 0.76 | | | |
| | Flower abundance | 0.0009 | 0.0006 | 14 | 1.52 | 0.15 | | | |
| | Management strategy | | | 2 | 0.07 | 0.94 | 9.67 (0.99, 18.36) | 11.21 (2.51, 19.92) | 12.13 (3.75, 20.51) |

| Response variable | Predictor | Standardized coefficient | Coefficient standard error | Degrees of freedom | $X^2$ | p-value | Continuous stocking coefficient (lower & upper confidence limits) | Low frequency rotation coefficient (lower & upper confidence limits) | High frequency rotation coefficient (lower & upper confidence limits) |
|---|---|---|---|---|---|---|---|---|---|
| Hymenoptera richness | Soil sand content | 0.02 | 0.38 | 14 | 0.04 | 0.96 | | | |
| | Flower abundance | -0.002 | 0.001 | 14 | -1.90 | 0.08 | | | |
| | Management strategy | | | 2 | 4.71 | **0.03** | 10.95 (-4.77, 26.66) | 3.56 (-12.20, 19.32) | 41.62 (26.45, 56.79) |
| Lepidoptera richness | Soil sand content | -0.01 | 0.12 | 14 | -1.06 | 0.31 | | | |
| | Flower abundance | 0.0009 | 0.0004 | 14 | 2.37 | **0.03** | | | |
| | Management strategy | | | 2 | 0.77 | 0.48 | 10.43 (5.07, 15.80) | 7.37 (1.98, 12.75) | 5.23 (0.05, 10.41) |

**Table S8:** Multivariate analysis of abundance investigating impacts of soil texture and management strategy on pollinator community composition. Terms significant at $\alpha$=0.05 are bolded.

| **Model term** | **$X^2$** | **Degrees of freedom** | **Residual degrees of freedom** | **p-value** |
|---|---|---|---|---|
| Intercept | | | 18 | |
| Management | 222.1 | 2 | 16 | **0.005** |
| Soil sand content | 181.2 | 1 | 15 | **0.001** |
| Flower abundance | 135.7 | 1 | 14 | **0.021** |

**Table S9:** Univariate likelihood ratio tests of impacts of soil texture and management strategy on pollinator taxa's abundance, from the community composition analysis. Terms significant at $\alpha$=0.05 are bolded.

| **Taxon** | **Predictor variable** | **Deviance** | **p-value** |
|---|---|---|---|
| *Abaeis nicippe* | Management | 3.46 | 1 |
| | Soil sand content | 2.32 | 1 |
| | Flower abundance | 1.05 | 1 |
| *Acmaeodera mixta* | Management | 5.76 | 0.96 |
| | Soil sand content | 0.45 | 1 |
| | Flower abundance | 0.89 | 1 |
| *Acmaeodera tubulus* | Management | 0.84 | 1 |
| | Soil sand content | 4.37 | 0.97 |
| | Flower abundance | 0.10 | 1 |
| *Apis mellifera* | Management | 0.43 | 1 |
| | Soil sand content | 0.31 | 1 |
| | Flower abundance | 5.96 | 0.90 |
| *Archytas apicifer* | Management | 1.05 | 1 |
| | Soil sand content | 0.87 | 1 |
| | Flower abundance | 0.14 | 1 |
| *Atalopedes huron* | Management | 4.02 | 1 |
| | Soil sand content | 0.09 | 1 |
| | Flower abundance | 0.56 | 1 |
| *Bombus pensylvanicus* | Management | 4.43 | 1 |
| | Soil sand content | 0.84 | 1 |
| | Flower abundance | 0.06 | 1 |
| Chloropidae spp. | Management | 2.89 | 1 |
| | Soil sand content | 0.02 | 1 |
| | Flower abundance | 0.37 | 1 |
| *Cryptorhopalum* spp. | Management | 1.45 | 1 |
| | Soil sand content | 12.65 | **0.02** |
| | Flower abundance | 0.83 | 1 |
| *Danaus plexippus* | Management | 5.39 | 0.98 |
| | Soil sand content | 1.17 | 1 |
| | Flower abundance | 3.04 | 1 |
| *Epicauta callosa* | Management | 3.50 | 1 |
| | Soil sand content | 1.14 | 1 |
| | Flower abundance | 0 | 1 |
| *Eristalis stipator* | Management | 5.76 | 0.96 |
| | Soil sand content | 0.19 | 1 |

| Taxon | Predictor variable | Deviance | p-value |
|---|---|---|---|
| | Flower abundance | 0.72 | 1 |
| *Lasioglossum disparile* | Management | 0.76 | 1 |
| | Soil sand content | 0.51 | 1 |
| | Flower abundance | 2.82 | 1 |
| *Mordellistena cervicalis* | Management | 3.07 | 1 |
| | Soil sand content | 0.06 | 1 |
| | Flower abundance | 0.51 | 1 |
| *Nathalis iole* | Management | 0.24 | 1 |
| | Soil sand content | 0.93 | 1 |
| | Flower abundance | 0.06 | 1 |
| *Odontocorynus salebrosus* | Management | 3.07 | 1 |
| | Soil sand content | 11.81 | **0.04** |
| | Flower abundance | 0.65 | 1 |
| *Polistes bellicosus* | Management | 4.61 | 1 |
| | Soil sand content | 0.03 | 1 |
| | Flower abundance | 2.41 | 1 |
| Sciaridae spp. | Management | 6.93 | 0.78 |
| | Soil sand content | 1.26 | 1 |
| | Flower abundance | 0.52 | 1 |
| *Sennius cruentatus* | Management | 2.51 | 1 |
| | Soil sand content | 0.29 | 1 |
| | Flower abundance | 0.29 | 1 |
| *Toxomerus marginatus* | Management | 1.17 | 1 |
| | Soil sand content | 1.79 | 1 |
| | Flower abundance | 0.01 | 1 |
| *Xylocopa virginica* | Management | 0.23 | 1 |
| | Soil sand content | 1.38 | 1 |
| | Flower abundance | 1.42 | 1 |
| *Andrena crawfordi* | Management | 1.40 | 1 |
| | Soil sand content | 0.05 | 1 |
| | Flower abundance | 0.18 | 1 |
| *Archytas californiae* | Management | 3.52 | 1 |
| | Soil sand content | 0.87 | 1 |
| | Flower abundance | 0.06 | 1 |
| *Colias eurytheme* | Management | 1.05 | 1 |
| | Soil sand content | 1.39 | 1 |
| | Flower abundance | 0.20 | 1 |

| Taxon | Predictor variable | Deviance | p-value |
|---|---|---|---|
| *Colletes birkmanni* | Management | 0.12 | 1 |
| | Soil sand content | 0.07 | 1 |
| | Flower abundance | 1.70 | 1 |
| *Epicauta ferruginea* | Management | 2.96 | 1 |
| | Soil sand content | 2.41 | 1 |
| | Flower abundance | 1.62 | 1 |
| *Epicauta pensylvanica* | Management | 0.68 | 1 |
| | Soil sand content | 4.05 | 1 |
| | Flower abundance | 1.75 | 1 |
| *Euptoieta claudia* | Management | 0.85 | 1 |
| | Soil sand content | 0.60 | 1 |
| | Flower abundance | 0.86 | 1 |
| Geron spp. | Management | 1.72 | 1 |
| | Soil sand content | 1.50 | 1 |
| | Flower abundance | 0.19 | 1 |
| *Junonia coenia* | Management | 1.42 | 1 |
| | Soil sand content | 0.47 | 1 |
| | Flower abundance | 1.16 | 1 |
| *Lasioglossum* sp. TX-03 | Management | 4.30 | 1 |
| | Soil sand content | 3.60 | 1 |
| | Flower abundance | 0.97 | 1 |
| *Lucilia illustris* | Management | 3.87 | 1 |
| | Soil sand content | 0.02 | 1 |
| | Flower abundance | 0.02 | 1 |
| *Polistes fuscatus* | Management | 0.04 | 1 |
| | Soil sand content | 5.38 | 0.90 |
| | Flower abundance | 11.49 | 1 |
| Sarcophaginae spp. | Management | 5.33 | 0.99 |
| | Soil sand content | 0.85 | 1 |
| | Flower abundance | 2.50 | 1 |
| *Sepsis* spp. | Management | 3.83 | 1 |
| | Soil sand content | 0.20 | 1 |
| | Flower abundance | 1.07 | 1 |
| *Stomoxys calcitrans* | Management | 2.94 | 1 |
| | Soil sand content | 0.12 | 1 |
| | Flower abundance | 6.04 | 0.89 |
| *Strymon melinus* | Management | 5.29 | 0.99 |
| | Soil sand content | 0.98 | 1 |
| | Flower abundance | 0.09 | 1 |

| Taxon | Predictor variable | Deviance | p-value |
|---|---|---|---|
| *Systoechus vulgaris* | Management | 1.73 | 1 |
| | Soil sand content | 0.06 | 1 |
| | Flower abundance | 0.18 | 1 |
| *Typocerus octonotatus* | Management | 5.61 | 0.97 |
| | Soil sand content | 0.39 | 1 |
| | Flower abundance | 0.05 | 1 |
| Anthomyiidae spp. | Management | 1.95 | 1 |
| | Soil sand content | 2.34 | 1 |
| | Flower abundance | 0.65 | 1 |
| *Coccinella septempunctata* | Management | 0.12 | 1 |
| | Soil sand content | 1.01 | 1 |
| | Flower abundance | 0.84 | 1 |
| *Dilophus* spp. | Management | 1.63 | 1 |
| | Soil sand content | 1.24 | 1 |
| | Flower abundance | 0.23 | 1 |
| *Entypus fulvicornis* | Management | 1.63 | 1 |
| | Soil sand content | 2.54 | 1 |
| | Flower abundance | 0.25 | 1 |
| *Eupeodes volucris* | Management | 5.11 | 0.99 |
| | Soil sand content | 2.21 | 1 |
| | Flower abundance | 0.52 | 1 |
| *Hippodamia convergens* | Management | 1.31 | 1 |
| | Soil sand content | 0.05 | 1 |
| | Flower abundance | 0.45 | 1 |
| *Lasioglossum hudsoniellum* | Management | 2.31 | 1 |
| | Soil sand content | 0.01 | 1 |
| | Flower abundance | 0.59 | 1 |
| *Lasioglossum semicaeruleum* | Management | 2.77 | 1 |
| | Soil sand content | 3.82 | 1 |
| | Flower abundance | 0.10 | 1 |
| *Lasioglossum* sp. TX-16 | Management | 2.26 | 1 |
| | Soil sand content | 3.10 | 1 |
| | Flower abundance | 0.97 | 1 |
| *Lasioglossum* sp. TX-06 | Management | 2.20 | 1 |
| | Soil sand content | 1.16 | 1 |

| Taxon | Predictor variable | Deviance | p-value |
|---|---|---|---|
| | Flower abundance | 0.18 | 1 |
| *Melissodes coreopsis* | Management | 1.18 | 1 |
| | Soil sand content | 8.63 | 0.26 |
| | Flower abundance | 0.12 | 1 |
| *Polistes apachus* | Management | 0.11 | 1 |
| | Soil sand content | 2.42 | 1 |
| | Flower abundance | 0.89 | 1 |
| *Andrena nothoscordi* | Management | 1.62 | 1 |
| | Soil sand content | 3.45 | 1 |
| | Flower abundance | 3.43 | 1 |
| *Bombus griseocollis* | Management | 3.36 | 1 |
| | Soil sand content | 0.18 | 1 |
| | Flower abundance | 1.41 | 1 |
| *Ceratina strenua* | Management | 1.68 | 1 |
| | Soil sand content | 10.47 | 0.10 |
| | Flower abundance | 0.001 | 1 |
| *Dilophus orbatus* | Management | 0.57 | 1 |
| | Soil sand content | 0.49 | 1 |
| | Flower abundance | 7.47 | 0.66 |
| *Erynnis horatius* | Management | 1.26 | 1 |
| | Soil sand content | 2.13 | 1 |
| | Flower abundance | 5.60 | 0.93 |
| *Lasioglossum* sp. TX-24 | Management | 1.74 | 1 |
| | Soil sand content | 2.27 | 1 |
| | Flower abundance | 0.29 | 1 |
| *Megachile brevis* | Management | 5.61 | 0.97 |
| | Soil sand content | 0.40 | 1 |
| | Flower abundance | 0.45 | 1 |
| *Myzinum quinquecinctum* | Management | 3.46 | 1 |
| | Soil sand content | 0.03 | 1 |
| | Flower abundance | 5.44 | 0.95 |
| *Trigonorhinus limbatus-tomentosus* | Management | 2.13 | 1 |
| | Soil sand content | 1.71 | 1 |
| | Flower abundance | 0.11 | 1 |
| *Diabrotica undecimpunctata* | Management | 6.86 | 0.83 |
| | Soil sand content | 0.66 | 1 |
| | Flower abundance | 0.09 | 1 |

| Taxon | Predictor variable | Deviance | p-value |
|---|---|---|---|
| *Nemognatha piazata bicolor* | Management | 5.19 | 0.99 |
| | Soil sand content | 1.22 | 1 |
| | Flower abundance | 0.39 | 1 |
| *Pontia protodice* | Management | 2.31 | 1 |
| | Soil sand content | 0.12 | 1 |
| | Flower abundance | 1.18 | 1 |
| *Acoloithus falsarius* | Management | 2.16 | 1 |
| | Soil sand content | 2.04 | 1 |
| | Flower abundance | 0.27 | 1 |
| *Megachile inimica* | Management | 1.40 | 1 |
| | Soil sand content | 0.40 | 1 |
| | Flower abundance | 1.98 | 1 |
| *Smicronyx* spp. | Management | 2.51 | 1 |
| | Soil sand content | 0.91 | 1 |
| | Flower abundance | 6.80 | 0.77 |
| *Megachile comata* | Management | 1.26 | 1 |
| | Soil sand content | 1.52 | 1 |
| | Flower abundance | 0.001 | 1 |
| *Acmaeodera ornatoides* | Management | 1.85 | 1 |
| | Soil sand content | 8.94 | 0.23 |
| | Flower abundance | 0.001 | 1 |
| *Andrena sitiliae* | Management | 1.86 | 1 |
| | Soil sand content | 6.51 | 0.70 |
| | Flower abundance | 0.001 | 1 |
| *Batyle suturalis* | Management | 1.26 | 1 |
| | Soil sand content | 0.01 | 1 |
| | Flower abundance | 2.06 | 1 |
| *Burnsius albescens-communus* | Management | 3.46 | 1 |
| | Soil sand content | 3.69 | 1 |
| | Flower abundance | 1.85 | 1 |
| *Diabrotica cristata* | Management | 1.85 | 1 |
| | Soil sand content | 9.35 | 0.20 |
| | Flower abundance | 0.001 | 1 |
| *Dielis plumipes* | Management | 3.91 | 1 |
| | Soil sand content | 3.77 | 1 |
| | Flower abundance | 4.48 | 0.99 |
| *Isodontia mexicana* | Management | 1.86 | 1 |

| Taxon | Predictor variable | Deviance | p-value |
|---|---|---|---|
| | Soil sand content | 1.39 | 1 |
| | Flower abundance | 6.01 | 0.90 |
| *Lithurgopsis gibbosa* | Management | 1.86 | 1 |
| | Soil sand content | 1.22 | 1 |
| | Flower abundance | 5.29 | 0.96 |
| *Poecilanthrx lucifer* | Management | 1.86 | 1 |
| | Soil sand content | 1.22 | 1 |
| | Flower abundance | 5.29 | 0.96 |
| *Psellidotus hieroglyphicus-johnsoni* | Management | 3.46 | 1 |
| | Soil sand content | 0 | 1 |
| | Flower abundance | 1.13 | 1 |
| *Villa* spp. | Management | 3.46 | 1 |
| | Soil sand content | 0 | 1 |
| | Flower abundance | 1.13 | 1 |
| *Systoechus solitus* | Management | 5.34 | 0.99 |
| | Soil sand content | 0.82 | 1 |
| | Flower abundance | 1.07 | 1 |
| *Euphoria kernii* | Management | 3.15 | 1 |
| | Soil sand content | 1.48 | 1 |
| | Flower abundance | 0.60 | 1 |
| *Halictus ligatus* | Management | 1.74 | 1 |
| | Soil sand content | 7.44 | 0.45 |
| | Flower abundance | 2.29 | 1 |
| *Melissodes boltoniae-tinctus* | Management | 1.85 | 1 |
| | Soil sand content | 2.41 | 1 |
| | Flower abundance | 5.27 | 0.96 |
| *Prionyx atratus* | Management | 1.26 | 1 |
| | Soil sand content | 4.88 | 0.94 |
| | Flower abundance | 2.16 | 1 |
| *Trichiotinus texanus* | Management | 1.86 | 1 |
| | Soil sand content | 2.10 | 1 |
| | Flower abundance | 3.51 | 1 |
| *Agapostemon angelicus* | Management | 1.26 | 1 |
| | Soil sand content | 2.60 | 1 |
| | Flower abundance | 0.13 | 1 |
| *Metrioidea popenoei* | Management | 1.85 | 1 |
| | Soil sand content | 0.58 | 1 |

| **Taxon** | **Predictor variable** | **Deviance** | **p-value** |
|---|---|---|---|
| | Flower abundance | 0.07 | 1 |
| *Polistes dorsalis* | Management | 1.26 | 1 |
| | Soil sand content | 0.22 | 1 |
| | Flower abundance | 0.11 | 1 |
| Scythrididae spp. | Management | 1.85 | 1 |
| | Soil sand content | 0.53 | 1 |
| | Flower abundance | 0.08 | 1 |

**Table S10:** Coefficients, test statistics, and post-hoc Tukey test p-values for models investigating effects of grazing management strategy and soil sand content on plant-pollinator network nestedness (weighted NODF) and network-level specialization ($H'_2$). Residual degrees of freedom are listed in the Intercept row for each model. Terms significant at $\alpha$=0.05 are bolded.

| **Response variable** | **Model term** | **Coefficient** | **Coefficient standard error** | **F** | **Degrees of freedom** | **p-value** | **Continuous vs. low frequency rotation** | **Continuous vs. high frequency rotation** | **Low vs. high frequency rotation** |
|---|---|---|---|---|---|---|---|---|---|
| Weighted NODF | Intercept | -3.14 | 1.01 | | 15 | | | | |
| | Low frequency rotation | -1.19 | 0.92 | 2.60 | 2 | 0.11 | 0.42 | 0.09 | 0.68 |
| | High frequency rotation | -1.89 | 0.83 | | | | | | |
| | Soil sand content | 0.04 | 0.03 | 2.21 | 1 | 0.16 | | | |
| $H'_2$ | Intercept | 7.46 | 1.99 | | 15 | | | | |
| | Low frequency rotation | 2.21 | 1.82 | 7.74 | 2 | **0.005** | 0.46 | **0.01** | 0.07 |
| | High frequency rotation | 6.20 | 1.64 | | | | | | |
| | Soil sand content | -0.08 | 0.06 | 1.79 | 1 | 0.20 | | | |

**Table S11:** This table is in an Excel file, and shows *c* and *z* values from hub analyses of networks at each site. Each sheet shows values for pollinators or plants at a specific site, or from a network that aggregated across all sites within a management strategy.

**Table S12:** Coefficients, test statistics, and post-hoc planned contrast p-values for models investigating effects of grazing management strategy pairs and distance on interaction dissimilarity (WN), interaction turnover (OS), and species turnover (ST). Planned contrasts asked whether dissimilarity (1) was greater among than within management strategies, and (2) differed among cross-strategy comparisons. Terms significant at α=0.05 are bolded.

| **Response variable** | **Predictor** | **Coefficient** | **Coefficient standard error** | **Degrees of freedom** | **$X^2$** | **p-value** | **Contrast 1** | **Contrast 2: Cont-LFR vs. Cont-HFR** | **Contrast 2: Cont-LFR vs. LFR-HFR** | **Contrast 2: Cont-HFR vs. LFR-HFR** |
|---|---|---|---|---|---|---|---|---|---|---|
| WN | Intercept | -1.87 | 0.78 | | | | | | | |
| | Comparison: LFR-LFR | 1.55 | 0.97 | 5 | 3.55 | 0.62 | 0.22 | 0.73 | 0.41 | 0.20 |
| | Comparison: HFR-HFR | 0.86 | 0.83 | | | | | | | |
| | Comparison: Cont-LFR | 0.95 | 0.62 | | | | | | | |
| | Comparison: Cont-HFR | 0.30 | 0.56 | | | | | | | |
| | Comparison: LFR-HFR | 0.68 | 0.74 | | | | | | | |
| | Distance (m) | 0.00008 | 0.00004 | 1 | 3.97 | **0.046** | | | | |
| OS | Intercept | -1.24 | 0.66 | | | | | | | |
| | Comparison: LFR-LFR | 0.73 | 0.85 | 5 | 6.76 | 0.24 | 0.28 | 0.74 | 0.84 | 0.62 |
| | Comparison: HFR-HFR | 0.89 | 0.71 | | | | | | | |
| | Comparison: Cont-LFR | 1.11 | 0.57 | | | | | | | |
| | Comparison: Cont-HFR | 0.22 | 0.52 | | | | | | | |
| | Comparison: LFR-HFR | 1.06 | 0.63 | | | | | | | |

| Response variable | Predictor | Coefficient | Coefficient standard error | Degrees of freedom | $X^2$ | p-value | Contrast 1 | Contrast 2: Cont-LFR vs. Cont-HFR | Contrast 2: Cont-LFR vs. LFR-HFR | Contrast 2: Cont-HFR vs. LFR-HFR |
|---|---|---|---|---|---|---|---|---|---|---|
| | Distance (m) | 0.00001 | 0.00003 | 1 | 0.15 | 0.70 | | | | |
| ST | Intercept | -1.19 | 0.79 | | | | | | | |
| | Comparison: LFR-LFR | 0.47 | 0.97 | 5 | 5.76 | 0.33 | 0.85 | 0.41 | 0.76 | 0.29 |
| | Comparison: HFR-HFR | 0.21 | 0.85 | | | | | | | |
| | Comparison: Cont-LFR | 0.12 | 0.61 | | | | | | | |
| | Comparison: Cont-HFR | 0.61 | 0.56 | | | | | | | |
| | Comparison: LFR-HFR | -0.24 | 0.75 | | | | | | | |
| | Distance (m) | 0.00007 | 0.00004 | 1 | 3.85 | **0.0498** | | | | |

**Figure S1:** Diagram depicting how each site was sampled. Flowers were identified and counted in six quadrats (green squares) placed along each of eight transects (dashed lines) radiating from the site's center. Soil texture (brown triangles) was averaged across samples taken at each site's center and the ends of four transects. Aerial netting was performed while walking clockwise along each of the eight transects.

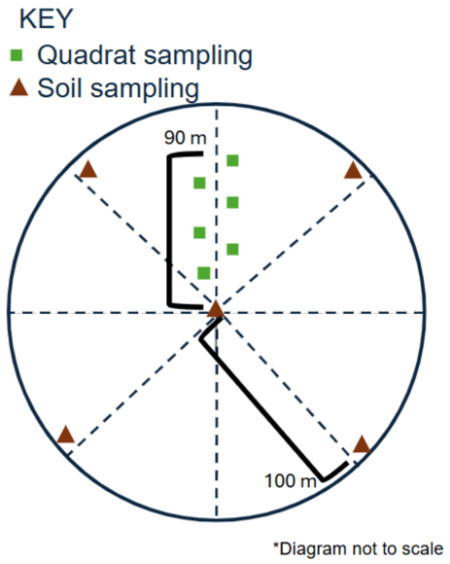

**Figure S2:** Soil texture diagram showing the percentage of clay, silt, and sand particles in soil from each site and sites' soil classifications. Cl indicates clay, Lo indicates loam, Sa indicates sandy, and Si indicates silt(y).

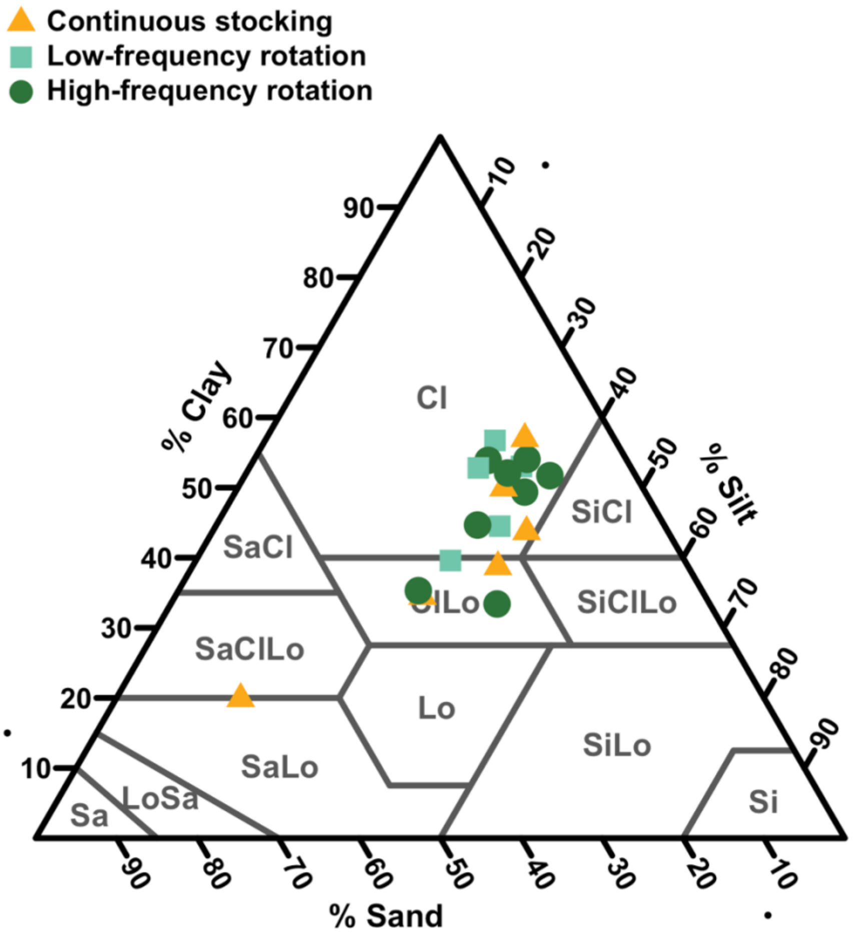

**Figure S3:** Impacts of (A) management and (B) soil texture on flowering plant richness. The line indicates the best fit line. A general linear model indicated that both terms impact flower richness (management $F_{2,15} = 9.02$, $p$=0.003; soil texture $F_{1,15} = 4.67$, $p$ = 0.047). A post-hoc Tukey test indicated that flower richness was higher with high frequency rotation than with continuous stocking ($p$ = 0.01) or low frequency rotation ($p$ = 0.005), but did not differ between continuous stocking and low frequency rotation ($p$ = 0.89).

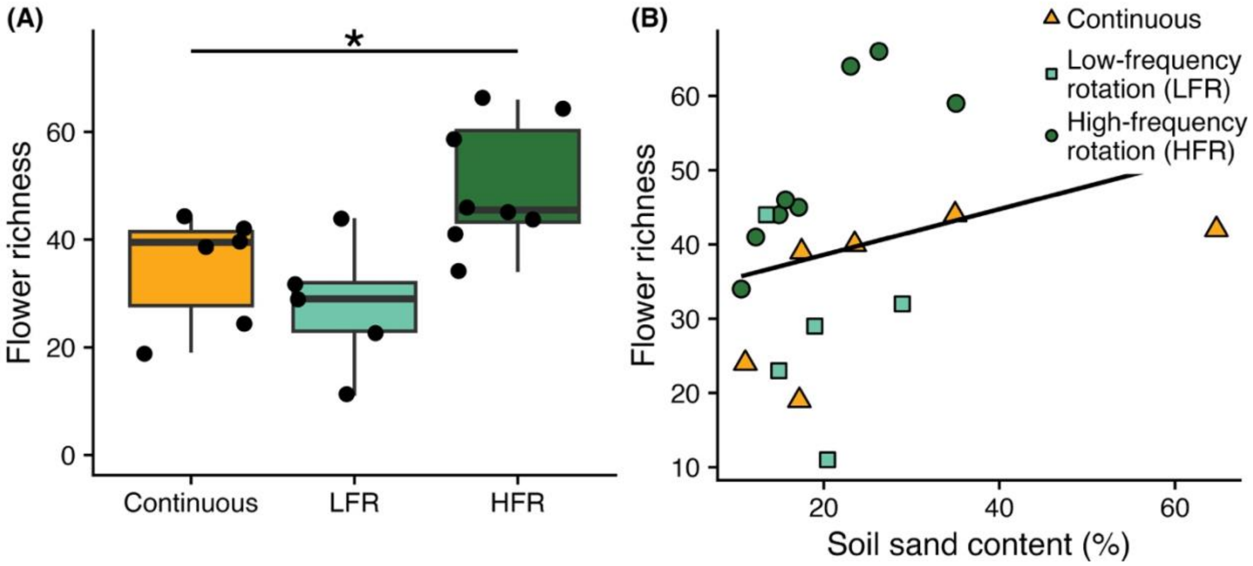

**Figure S4:** Impacts of flower abundance on (A) Coleoptera, (B) Diptera, (C) Hymenoptera, and (D) Lepidoptera abundance. The curves indicate the best fit curves.

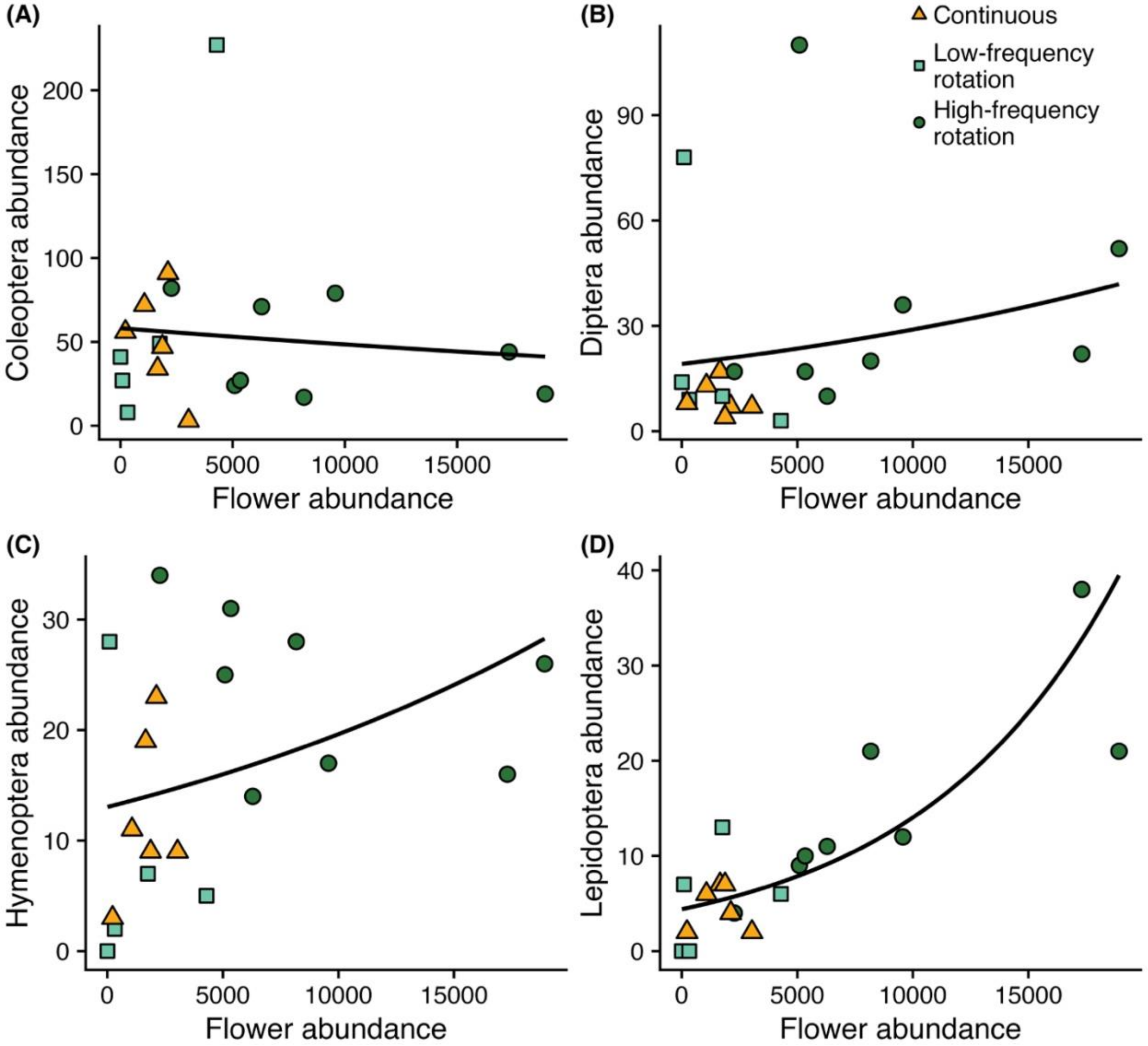

**Figure S5:** Impacts of flower abundance on (A) Coleoptera, (B) Diptera, (C) Hymenoptera, and (D) Lepidoptera richness. Richness was estimated using the iNEXT R package. The lines indicate the best fit lines.

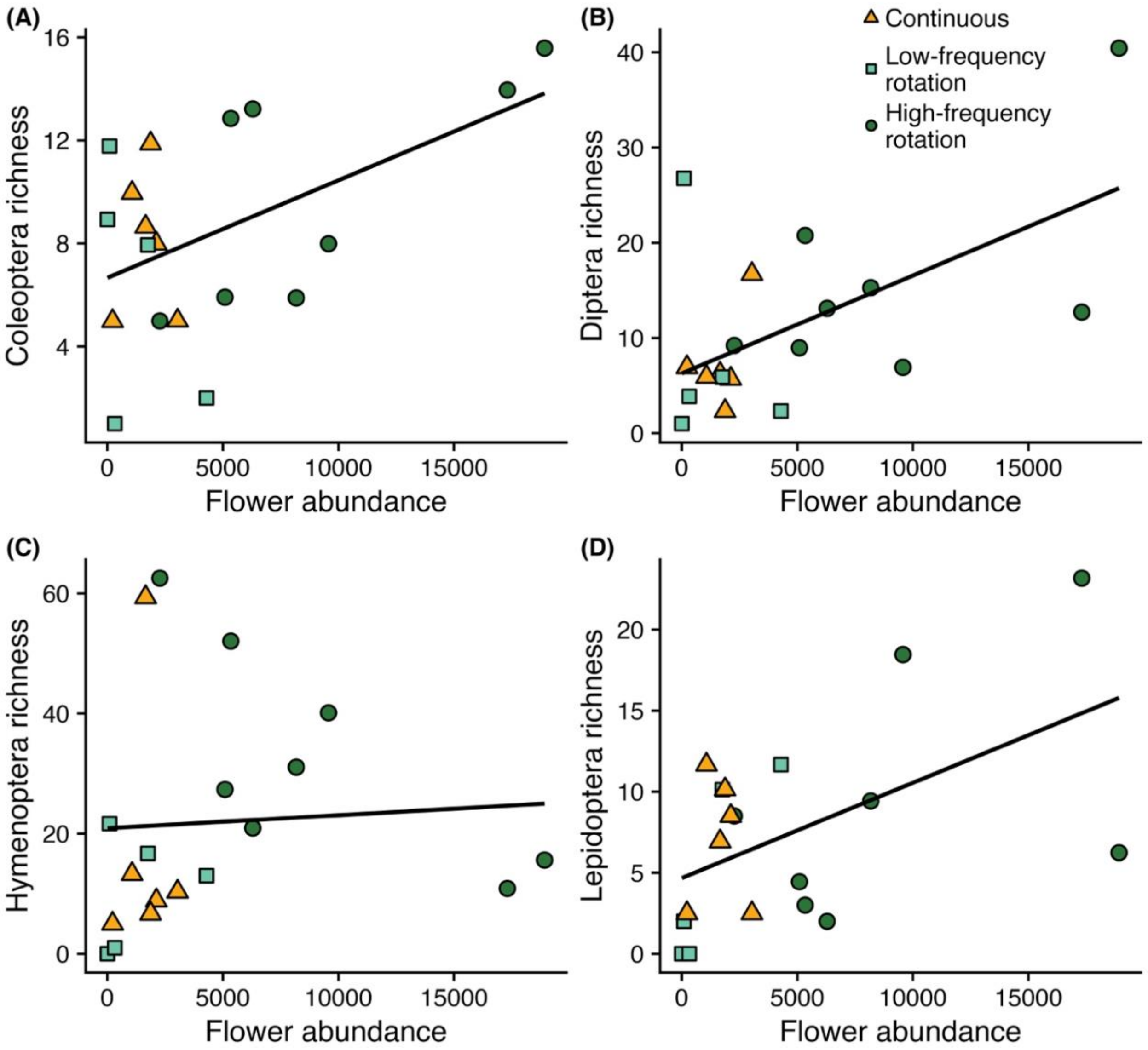
